# Atomic model of Vesicular Stomatitis Virus and Mechanism of Assembly

**DOI:** 10.1101/2022.05.02.490251

**Authors:** Kang Zhou, Zhu Si, Peng Ge, Jun Tsao, Ming Luo, Z. Hong Zhou

**Affiliations:** Department of Microbiology, Immunology & Molecular Genetics, University of California, Los Angeles (UCLA), Los Angeles, CA 90095, USA; California NanoSystems Institute, UCLA, Los Angeles, CA 90095, USA; School of Life Science, University of Science and Technology of China, Hefei, Anhui 230026, P.R. China; Department of Microbiology, University of Alabama at Birmingham, Birmingham, Al 35294, USA; The Department of Chemistry, Georgia State University, Atlanta, GA 30303, USA

**Keywords:** Vesicular stomatitis virus, Helical nucleocapsid, cryoEM, cryoET, Matrix protein M, Nucleocapsid protein N, Glycoprotein G

## Abstract

Like other negative-strand RNA viruses (NSVs) such as influenza and rabies, vesicular stomatitis virus (VSV) has a three-layered organization: a layer of matrix protein (M) resides between the membrane envelope, studded by glycoprotein (G), and the nucleocapsid, composed of the nucleocapsid protein (N) and the encapsidated genomic RNA. Lack of *in situ* atomic structures of these viral components has limited our understanding of the virion assembly mechanism. Here, by cryoEM and sub-particle reconstruction, we have determined the *in situ* structures of M and N inside VSV at 3.47 Å resolution. In the virion, N and M have a stoichiometry of 1:2. The *in situ* structures of both N and M differ from their crystal structures in their N-terminal segments and oligomerization loops. N-RNA, N-N, and N-M-M interactions govern the formation of the capsid. A double layer of M contributes to packaging of the helical nucleocapsid: the inner M (IM) joins neighboring turns of the N helix, while the outer M (OM) contacts G and the membrane envelope. The pseudo-crystalline organization of G is further mapped by cryoET. The mechanism of VSV assembly is delineated by the network interactions of these viral components.

## Introduction

Negative-strand RNA viruses (NSVs) constituting the *Haploviricotina* subphylum of the *Negarnaviricota* phylum include some of the most devastating human pathogens, such as rabies virus (RABV) (*Rhabdoviridae*), measles virus (MeV) (*Paramyxoviridae*), influenza virus (*Orthomyxoviridae*), and Marburg and Ebola viruses (*Filoviridae*). Vesicular stomatitis virus (VSV) is an enveloped, bullet-shaped rhabdovirus, which is closely related to the highly contagious and historically significant RABV^1,2^. VSV is the prototypical NSV and has long been used as a model for RABV and other NSVs. It has also been widely used for engineering pseudotypes—VSV-like particles carrying effector molecules—as both vaccines (including those against SARS-CoV-2, the virus responsible for the COVID-19 pandemic^3,4^) and anti-cancer agents^5^. Attenuated VSV strains are non-toxic to normal tissues, as they effectively trigger an interferon-mediated anti-viral response against themselves^6^. Vaccines based on pseudo-type VSV carrying foreign surface proteins or recombinant VSV expressing foreign viral genes have been developed as vaccine candidates for COVID-19 (ref. ^7^), human immuno-deficiency virus (HIV)^8,9^, “avian flu”^10^, measles^11^, Ebola virus^12^ and Marburg virus^12,13^. Pseudotypes of VSV that carry HIV receptors selectively target and kill HIV-1 infected cells and control HIV-1 infection^14,15^. The atomic details of the molecular interactions governing VSV assembly would benefit all these therapeutic and prophylactic endeavors.

All members of *Rhabdoviridae* encode three structural proteins: nucleocapsid protein (N), matrix protein (M) and glycoprotein (G). In the virion, these proteins are organized into a distinctive bullet shape. M is sandwiched between a G-containing membrane envelope and a nucleocapsid, composed of N and a single-stranded genomic RNA. The crystal structures have been reported for the C-terminal core domain of M (*i*.*e*., M^t^)^16^, N^17^ and the two forms of the ectodomain of G^18,19^. Previous cryo electron microscopy (cryoEM) structures of VSV from helical reconstruction at 13-Å resolution have led to a model of 3’ to 5’ RNA-guided assembly of the nucleocapsid from the tip of the bullet to the trunk, forming a bullet-shaped viral particle^20^. However, without *in situ* structures of M and N in the virion at a near-atomic resolution, the precise arrangement of M, N and G and their interactions governing the virion assembly remain elusive, limiting our understanding of VSV assembly and impeding rational engineering efforts of VSV pseudotypes.

In this study, we determined the *in situ* structures of M and N inside VSV at 3.47 Å resolution by cryoEM and sub-particle reconstruction. The structure shows that each N interacts with a pair of M subunits in the virion, not the previously reported single M. Cryo electron tomography (cryoET) and subtomogram averaging further establish the pseudo-crystalline organization of G with predominantly hexagonal and occasionally pentagonal tiles on the membrane envelope. The interactions between N and M, as well as matching distributions of M and G, suggest a mechanism of VSV assembly.

## Results

### *In situ* structures of M, N and encapsidated RNA resolved at near atomic resolution by helical sub-particle reconstruction

The large variable range of subunits/turn (35.5-41.5 subunits/turn, though previous studies have shown mostly 37.5 subunits^20^) has hitherto presented challenges in reconstructing the VSV trunk to a resolution needed for atomic modeling using the conventional helical reconstruction method. Here, we were able to determine the *in situ* structure of the VSV trunk to 3.47 Å resolution (Fig. 1 and Supplementary Video 1) by subjecting particles to fine 3D classification based on subunits/turn and then treating small regions in the major class (38.5 subunits/turn) independently, a novel approach which we termed helical sub-particle reconstruction (see Method for details). The structure reveals the 1:2 ratio between N and M and resolves amino acid side chains and RNA bases, both of which were used to build the atomic models (Fig. 1c and Supplementary Video 1).

**Fig. 1.**
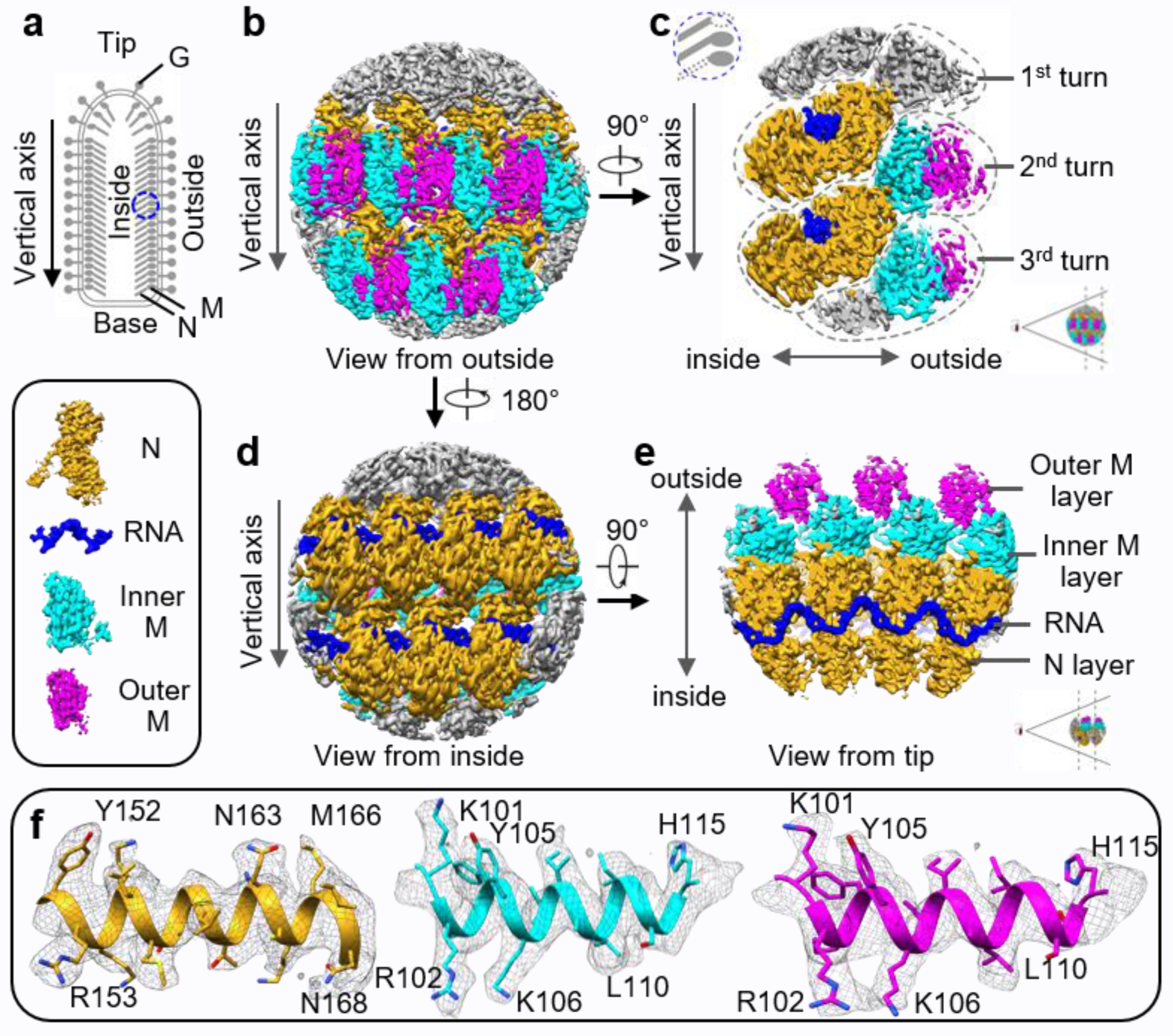
Sub-particle reconstruction of VSV trunk at near-atomic resolution. **a**, Schematic diagram of the VSV virion. Blue circle marks the region used for sub-particle reconstruction. **b** and **d**, CryoEM density maps of partial VSV nucleocapsid shown in two opposite views. Only intact N, intact M and RNA are colored for clarity; other densities are gray. N subunits are colored in goldenrod, inner M in cyan, outer M in magenta and RNA in blue (inset). **c** and **e**, Cross sections of cryoEM density maps viewed from side and tip. **f**, CryoEM density maps (mesh) of α-helices from three kinds of subunits.

We were able to fit an atomic model of 19 protein subunits, comprising 7 N and 12 M, and two long segments of single stranded RNA (33 and 35 nucleotides in length, respectively) into our density map obtained by sub-particle reconstruction (Fig. 2b, c and Supplementary Video 2). Based on the extent of interactions between subunits, we defined an asymmetric unit to contain an N subunit and two M subunits (Fig. 2e). The atomic model contains 19 subunits arranged across three radial layers (from inner to outer: N layer, inner M (IM) layer, and outer M (OM) layer in one intact and two partial helical turns of the asymmetric units along the helical axis (Fig. 2c). Within the same turn, each IM subunit interacts with two N subunits, one N from the same asymmetric unit and another from a neighboring asymmetric unit (Fig. 2f-h). This IM subunit is also inlaid between two OM subunits, one OM from the same asymmetric unit and another from a neighboring asymmetric unit (Fig. 2h). Each IM subunit is placed between N subunits from two successive (upper-lower) turns (Fig. 2d), but the IM subunit in lower turn has no significant interactions with the N subunit from upper turn. In addition to the close lateral interactions between two adjacent N subunits in the same turn, the single-stranded RNA threads through the cleft of N subunits (Fig. 2h). Each N subunit accommodates 9 nucleotides of RNA (Fig. 2e).

**Fig. 2.**
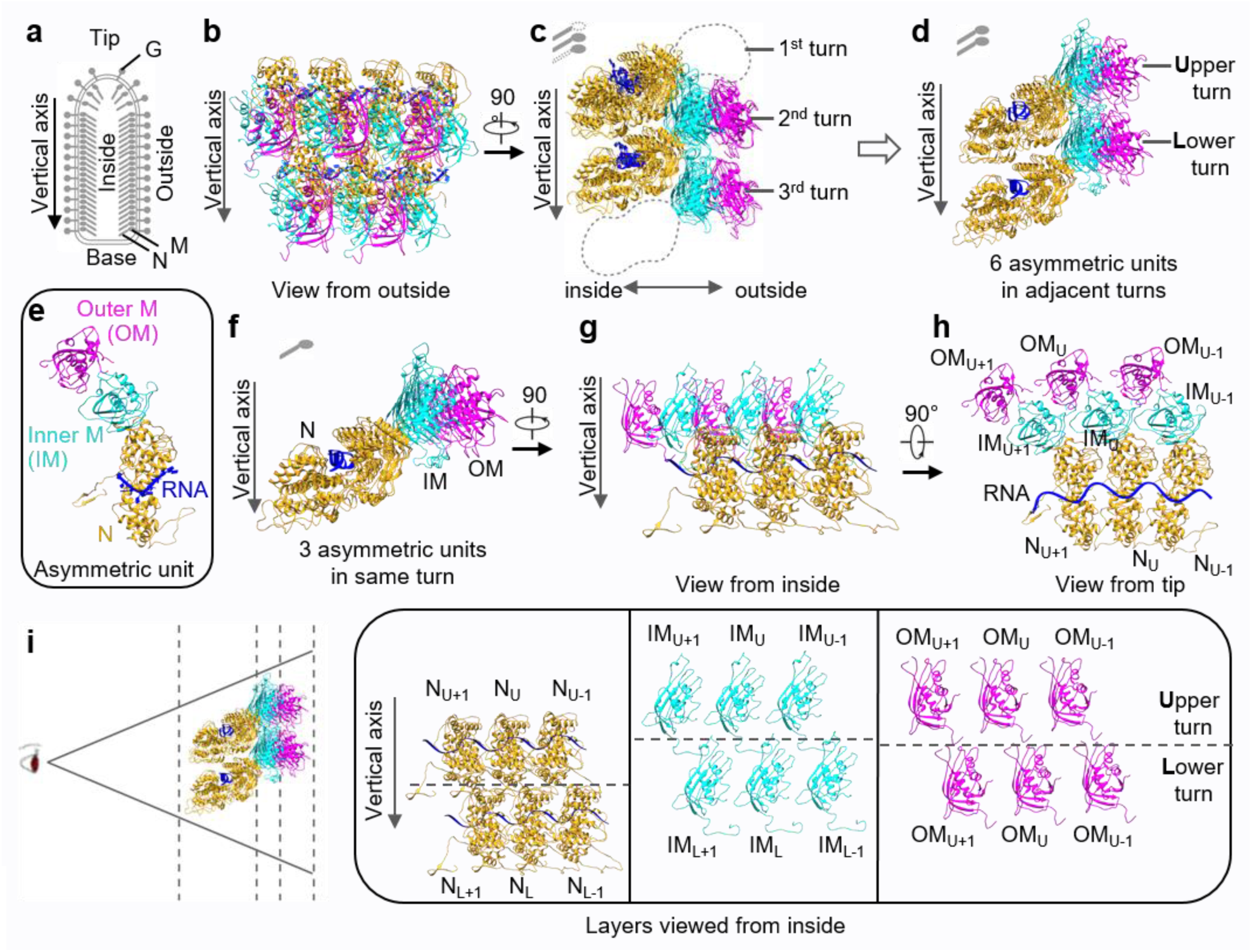
Atomic model of the partial VSV nucleocapsid. **a**, Schematic diagram of the VSV virion. **b, c**, Two orthogonal views of the atomic model, which contains 7 N (goldenrod), 7 inner M (cyan) and 5 outer M (magenta). Only intact N, intact M and RNA are built and colored, as in the cryo-EM map. **d**, Six asymmetric units from two turns derived from (**c**). **e**, One asymmetric unit, including 1N, 1 IM and 1 OM, together with 9 RNA nucleotides. **f-h**, Three asymmetric units from the same turn shown in three orthometric views. **i**, Layer expansion of (**d**) from the central axis of the virion. Each subunit is labeled based on its lateral relationship (+1 or -1, along the left-handed helix) and vertical position (Upper (U) and Lower (L) turns).

### Comparison between *in situ* and crystal structures of N

The *in situ* atomic model of the full-length N (Fig. 3a) exhibits significant differences from the crystal structure (PDB: 2GCI)^17^ of recombinantly-expressed N. The crystal structure comprises 10 subunits in one turn, or a decamer, while each turn in the *in situ* VSV trunk ranges from 35.5 to 41.5 subunits. When the *in situ* and crystal structures of single N subunit are compared, the main body and RNA-binding pattern remain the same between two structures, but there are significant differences in the N-terminal arm (Ser2 – Val25, termed N-arm) and an extended loop (Leu344 – Tyr355, termed C-loop) (Fig. 3a). Compared to those in the crystal structure (blue), in *in situ* N (red), the N-arm rotates downwards (towards the base) by 30 degrees, and the C-loop rotates outwards (towards the outside of the capsid) by 13 degrees (Fig. 3a). Superposition of the crystal and *in situ* structures of three adjacent N subunits was then performed by aligning only the middle N subunits (N_U_) (Fig. 3b) while allowing the adjacent subunits to be not aligned. Interestingly, a stable interlocking anchor is found in the interactions among the three N subunits in both the crystal and the *in situ* structures (Fig. 3b and inset). The N-arm from the first N subunit (N_U-1_) forms a β-hairpin and extends along the edge of N_U_. On the opposite side, the C-loop from the third N subunit (N_U+1_) runs over the C-domain of N_U_ and doubles back. From the outside view of the nucleocapsid, the N-arm from the N_U-1_ subunit and the C-loop from the N_U+1_ subunit form a “handhold,” which holds in place the N_U_ subunit (Fig. 3b, left); the interactions and relative position between the “handhold” and the N_U_ are constant across the turn (Fig. 3b inset). In this architecture, one N subunit connects its N-arm and C-loop to the N that are two subunits away on either side (N_U-2_, N_U+2_). Adjusting the conformation of the N-arm and C-loop allows N to adopt to different subunit numbers per turn with different diameters. In the *in situ* structure, the N-arm and C-loop pivot outwards, which makes the ring larger to accommodate more subunits per turn.

**Fig. 3.**
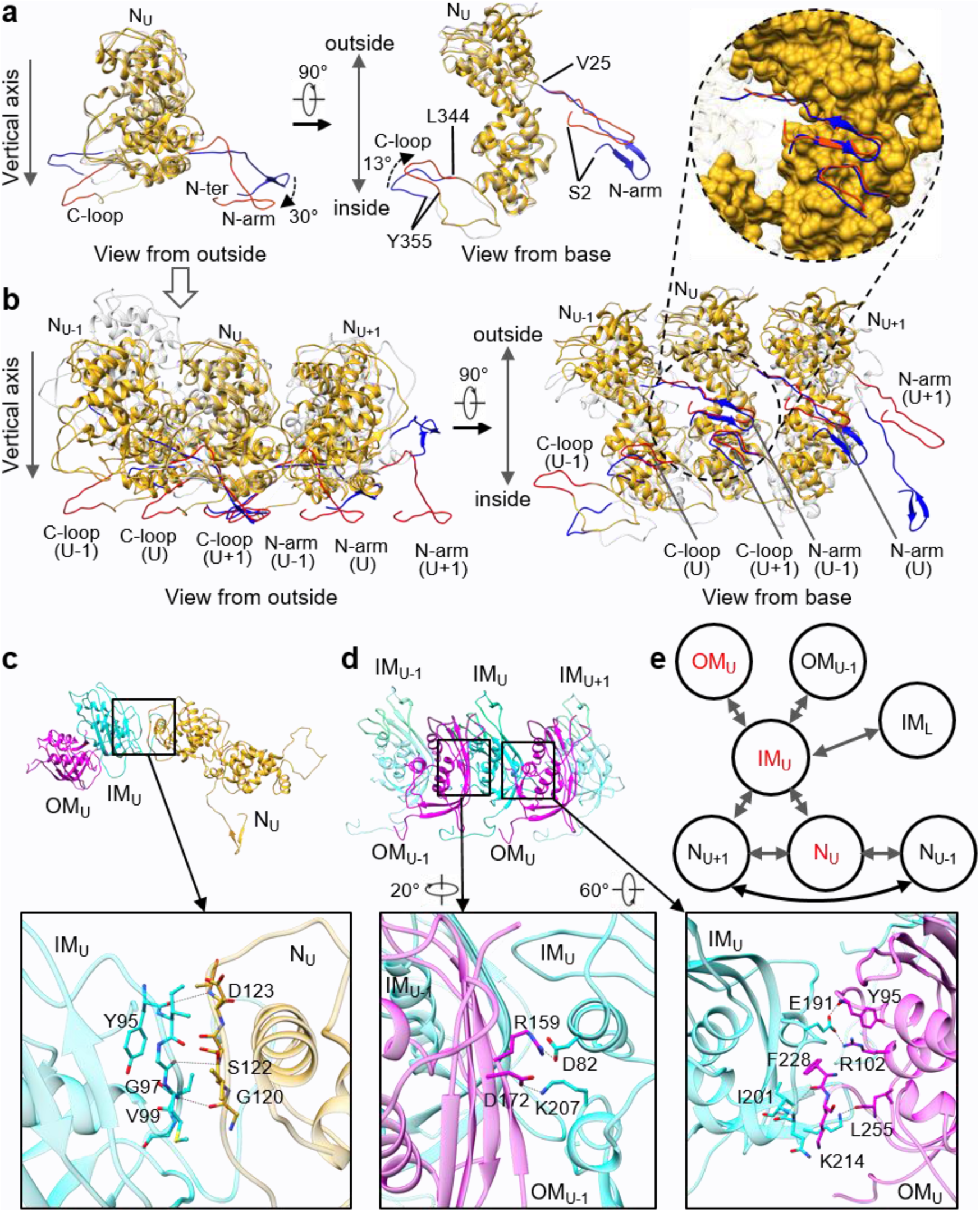
Analysis and illustration of subunits interactions. **a**, Superposition of the *in situ* and crystal structures of a single N subunit. The *in situ* structure is colored in goldenrod except for the N-arm and C-loop, in red, while the crystal structure is colored in transparent gray except for the N-arm and C-loop, in blue. **b**, Superposition of three N subunits extended from (**a**). Only the middle N subunits are aligned. Inset: N_U_ is shown as surface contour. **c**, Interactions between IM and N from the same asymmetric unit. **d**, Interactions between IM and two OMs, one OM from the same asymmetric unit and another one from the adjacent asymmetric unit. **e**, The subunit interaction network in the VSV nucleocapsid, in which IM plays a key mediating role. As labeled, subunits N_U_, IM_U_ and OM_U_ belong to the same asymmetric unit.

### Comparison between *in situ* and crystal structures of M and between IM and OM

The final sub-particle reconstruction clearly shows that there is a double layer of M surrounding the helical nucleocapsid in the virion (Figs. 1c and 2c). The IM layer bridges between the N layer and the OM layer; there is no direct contact between N and OM (Fig. 2c, d, f). The distances between the lateral IM subunits, as well as between the lateral OM subunits, are too large to establish significant IM-IM or OM-OM interactions (Fig. 2h, i).

The crystal structure of M^21^ (PDB: 2W2R) was fitted into the corresponding EM densities. The main body of the crystal structure fit well, but an N-terminal segment of 12 residues (Leu41 – Asp52) could not fit. A segment of five residues (Leu53 – Asp57) is missing in the crystal structure of M (Extended Data Fig. 1a); however, the distance between Asp52 and Ser58 in the EM density is 46.6 Å (Extended Data Fig. 1b), too large to be spanned by five amino acid residues even when the main-chain is fully extended. Thus, the 12-residue segment and the main body next to each other cannot belong to the same M subunit in the *in situ* structure. In the crystal structure of M, the 12-residue segment could be assigned to another M molecule in the neighboring asymmetric unit^21^. After examining the EM densities and locations of M subunits, we focused on a pair of IM subunits, one from the upper helical turn (IM_U_) and the other from the lower turn (IM_L_) (Fig. 2i and Extended Data Fig. 1b). The distance between Asp52 (next to IM_L_) and Ser58 (IM_U_) is only 10.5 Å (Extended Data Fig. 1b), which is much shorter than the previously mentioned distance of 46.6 Å and within the range for inserting five residues. Based on this observation, the 12-residue segment (Leu41 – Asp52, beside IM_L_) was assigned to IM_U_, and five residues were added to that segment after Asp52 (Leu53 – Asp57) (Extended Data Fig. 1a, b). We also observed that the 12-residue segment interacts with IM_L_ (Extended Data Fig. 1b). This completes an *in situ* atomic model of IM from residue Leu41 to the last residue at the C terminal end (Extended Data Fig. 1a), but the N-terminal 40 residues (Met1 – Pro40) remain disordered.

Superposition of IM and OM reveals three differences between the two molecules (Extended Data Fig. 1c). First, the 12-residue segment (Leu41 – Asp50) is missing in OM. Second, residues Val122 – Asn127 in OM are also missing, and the segment from Asn194 to Ser199 in OM has a conformation different from that in IM (Extended Data Fig. 1c). All three conformational changes of OM are located at its outward-facing regions, far away from N and IM layers but next to the viral envelope. Third, IM and OM are oriented completely differently inside the virion (Fig. 2i). These changes suggest that IM and OM play different roles in capsid assembly.

### Subunit interactions governing virion assembly

To understand VSV virion assembly, we used the PISA software^22^ to calculate the Gibbs free energy and the interface area between subunits in order to quantify inter-subunit interactions in the nucleocapsid (Supplementary Table 1). The lateral interactions of adjacent N subunits are the strongest, while those of N_U-1_-N_U+1_ subunits are weaker but still significant (Fig. 3b and Supplementary Table 1, rows in blue). Remarkably, although N subunits from successive turns have a buried interface, there are no significant vertical interactions between the N subunits (Fig. 2d and Supplementary Table 1, third and fourth rows).

Despite its location between two successive turns of N subunits (Fig. 2d), the IM subunit interacts significantly only with two N subunits in the same turn, not the N in any other turns (Fig. 2h, i and Supplementary Table 1, rows in orange). These interactions occur on the inner side of the IM. The residues D123/S122/G120 from N_U_ and Y95/G97/V99 from IM_U_ comprise the main interactions between the two (Fig. 3c). There is a buried interface between IM_L_ and N_U_, but there are no strong interactions between them (Fig. 2i and Supplementary Table 1, fifth row). On its outer side, the IM subunit interacts with two independent OM subunits (Fig. 2h). One OM subunit is in the same asymmetric unit as the IM subunit and binds to that IM subunit more strongly than the other OM subunit from the neighboring asymmetric unit (Fig. 2h and Supplementary Table 1, rows in green); residues involved include E191/F228/I201/D82/K207 from IM_U_, Y95/R102/L255/K214 from OM_U_ and R159/D172 from OM_U-1_ (Fig. 3d).

In general, the interactions among subunits can be classified into three types according to their directions: lateral, radial and vertical (Fig. 3e). The lateral interactions are mainly at the N-N interfaces (Fig. 3b), including those between N_U_ and N_U+1_ and between N_U-1_ and N_U+1_. The lateral interactions are enhanced by the single-stranded RNA woven through the N subunits (Fig. 2h). The radial interactions are mediated by IM subunits: one IM subunit interacts with two N subunit inwards and with two OM subunits outwards (Fig. 2h). The main vertical interactions are between the N-terminal segment (Leu41 – Met48) of IM_U_ and the main body of IM_L_ (Fig. 2i and Extended Data Fig. 1b), and those interactions determine the distance between successive turns (Extended Data Fig. 1b). These three types of interactions govern the structure of the virion but do not explain the attachments of G.

### Super-complexes of G trimers on the virion surface

In order to explore the overall organization of M, N and G and the interactions between them, we recorded 30 cryoET tilt series for tomographic analysis. The reconstructed tomograms (Fig. 4a-d and Supplementary Video 3), especially those after denoising (Fig. 4b, d), resolve the 3D organizations of VSV virions. The regularly patterned M and N proteins inside the virion are clearly visible in the tomograms, as are the G proteins that exist as trimers on the viral membrane envelope (Fig. 4c, d). Most G proteins on the envelope are distributed across the virion except for on the base, which has few if any G proteins. The vast majority of the G trimers were already in the taller and thinner postfusion conformation^18^ under the neutral pH condition used during our experiment (pH=7.4); some shorter and wider G trimers in the prefusion conformation^19^ were also present (Fig. 4e-g). Remarkably, we observed hexagonal-shaped tiles in our reconstructed tomograms, as well as occasional pentagonal tiles in places such as the tip of the virion (Fig. 4h, i). These hexagon-hexagon or hexagon-pentagon tiles form local clusters of G trimers on the viral membrane envelope, suggesting that the G trimers follow a specific organization during virion assembly.

**Fig. 4.**
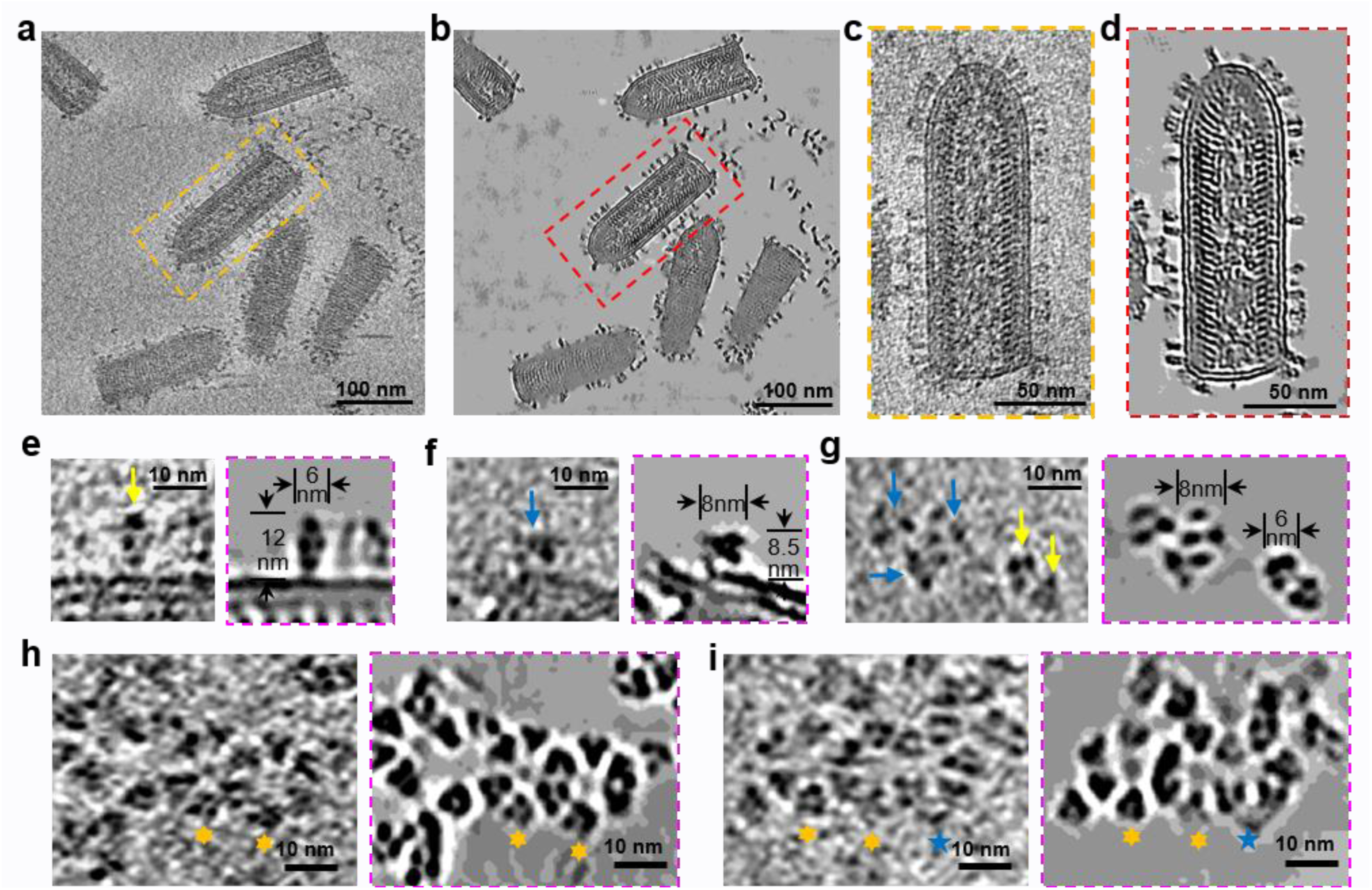
*In situ* structures of VSV G trimers and their hexagonal/pentagonal distribution super-complexes visualized by subtomogram averaging. **a, b**, An 8 Å-thick density slice from a reconstructed tomogram. (**b**) is the denoised image of (**a**). **c, d**, A representative VSV virion, corresponding to the one virion boxed in (**a, b**). (**d**) is the denoised image of (**c**). **e-g**, VSV G trimers in prefusion or postfusion conformation. (**e**) are side views of a representative G trimer in postfusion conformation; (**f**) are side views of a representative G trimer in prefusion conformation; (**g**) are top views of G trimers in both prefusion and postfusion conformation. Yellow arrows indicate postfusion conformation; blue arrows indicate prefusion conformation. Images with magenta dashed lines are the corresponding denoised images. **h, i**, Examples of hexagonal and pentagonal tiles of G trimers on the viral membrane. (**h**) shows two connected hexagonal tiles; (**i**) shows two connected hexagonal tiles joined to a pentagonal tile. Hexagonal tiles and pentagonal tiles are denoted by gold six-pointed stars and blue five-pointed stars, respectively. Images in magenta dashed boxes are the corresponding denoised images.

By averaging a total of 6030 subtomograms, an *in situ* structure of the G trimer was obtained at a resolution of 14.9 Å (Fig. 5a, first row, above the dashed line), which reveals three blurred density blobs around the central G trimer. The blobs, all similarly sized, appear to be trimeric and even contact a protomer (*i*.*e*., a G monomeric subunit) of the central G trimer (leftmost panel in the first row of Fig. 5a). The neighboring trimeric blobs are smeared due to the 3-fold symmetry of the central G trimer and the curvature of the virion. Subsequent symmetry relaxation and refinement yielded four different categories of G trimer super-complexes, which all contain a central G trimer but different numbers of neighboring G trimers (Fig. 5a, below the dashed line). Among the four types of super-complexes, the super-complex containing one neighboring G trimer has the largest number of subtomograms (42.8% of the total subtomograms), followed by ones with two neighboring (33.2%), none neighboring (15.2%) and three neighboring (8.8%) (Fig. 5a); the resolution of the corresponding averaged structures are 19.6 Å, 20.9 Å, 18.4 Å and 20.9 Å, respectively (Extended Data Fig. 2c). The orthogonal sectional views of the super-complexes clearly show that the central G trimer contacts its neighboring trimers (Fig. 5a, third column). The superposition of the super-complex density map with atomic models reveals possible contact sites at the N-terminus (residues 8–14) of each protomer (Extended Data Fig. 2d, e). These potential contacts suggest that the organization of G trimers in such super-complexes is not randomly formed by close packaging of individual G trimers.

**Fig. 5.**
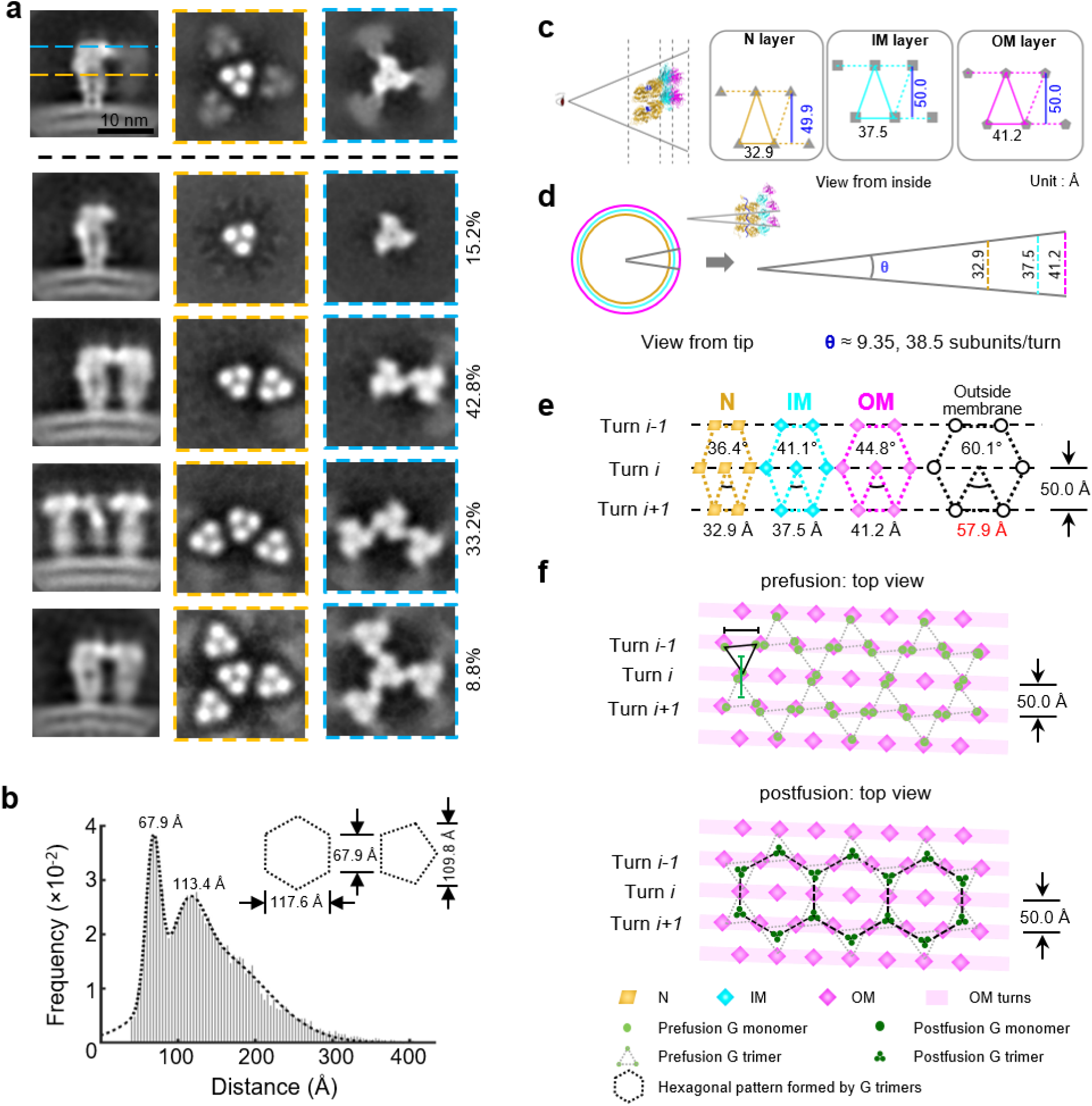
The super-complexes of G trimers visualized by subtomogram averaging and the relationship between the spatial organization of OM and G. **a**, Structures of G trimer before classification and G trimers classified by different numbers of neighboring G trimers (super-complexes), obtained by subtomogram averaging. Images above the black dashed line (first row) are the averaged density map of all subtomograms. Images below black dashed line are the averaged density maps of G trimer with 0 (second row), 1 (third row), 2 (fourth row) and 3 (fifth row) neighboring G trimers. Images in the first column are side views of the averages; images in the second and third column are orthogonal slice views; location is indicated by blue and golden dashed lines in the first image. **b**, Distribution of the distances of the first five nearest neighbors for each G trimer. The dashed curve shows the fitting of a three-term Gaussian model. The second peak (113.4 Å) is between the widths of hexagon (117.6 Å) and pentagon (109.8 Å), which have an identical side length (67.9 Å, the first peak), indicating a mixture of hexagonal and pentagonal tiles. **c, d**, Layer expansion of six asymmetric units in two turns; subunits are simplified to triangle (N), square (IM) and pentagon (OM). The measured distance between turns for each subunit is about 50.0 Å. Lateral distances between neighboring subunits in N layer, IM layer and OM layer are proportional to the radius of the three layers. **e**, Predicted side length of the hexagonal tiles of G trimers based on the lateral distances of neighboring N, IM and OM. **f**, Hexagonal arrangement of G trimers. The distances between OMs underneath the membrane match the distances between prefusion G trimers outside the membrane, as measured from the position of their fusion loops. In the postfusion conformation, the fusion loops in the G trimers are closer to the trimer axis.

Our direct observation of the hexagonal tiles is consistent with previous proposals that G proteins follow a regular arrangement on the viral membrane envelope due to the tendency of G to trimerize and interact through the endodomain of G with the underlying M lattice^20^. Statistically, the distances between neighboring G trimers in the super-complexes peaked sharply at 67.9 Å, corresponding to the side length of both hexagons and pentagons formed by the postfusion G trimers (Fig. 5b). Interestingly, we also observed a second peak at 113.4 Å, which is bigger than the distance of the diagonal in the postfusion pentagon (*i*.*e*., 109.8 Å) but smaller than the distance of the short diagonal (117.6 Å) in the postfusion hexagon (Fig. 5b). In addition, this second peak is flatter than the first peak, suggesting that it consists of an average of the two vertex distances of the pentagon and hexagon. These two statistical observations are consistent with our observed visualization of both hexagonal and pentagonal tiles among the G trimers in our postfusion structure.

In the crystal structure of the prefusion G trimers, the distance between the centers of adjacent trimers is 69 Å^19^ (Extended Data Fig. 3d). In the *in situ* structure of the G trimers in the postfusion conformation, we observed the distance between the centers of adjacent trimers to be 67.9 Å, which is very close to the distance between the centers of the prefusion G trimers. Additionally, the distance between the monomeric subunits of the crystal structure prefusion G trimers is 58 Å^19^ (Extended Data Fig. 3d). In order to calculate where the monomeric subunits would be given our *in situ* observations, we reviewed previous studies that have suggested that G interacts with M^23^. Thus, using *in situ* measurements of the arrangement of N, IM, and OM, given that the three helices formed by N, IM and OM are coaxial and the distances between neighboring subunits are proportional (Fig. 5c, d), we calculated the location of G. This calculation resulted in a distance of 57.9 Å between the hypothetical monomeric subunits (Fig. 5e), which is also quite close to the distance between the actual monomeric subunits of the prefusion G trimers in the crystal structure. Thus, the observed crystal structure distances, of 69 Å between G trimers and of 58 Å between G monomeric subunits, align with the observed and calculated *in situ* distances, of 67.9 Å between G trimers and of 57.9 Å between G monomeric subunits.

The alignment of these distances suggests that the three endodomains of a prefusion G trimer bind to three different OM subunits spanning two turns (Fig. 5f). Moreover, two monomeric subunits from two adjacent G trimers would occupy the same vertex of a hexagon, which suggests that the endodomains of these two subunits are positioned to contact the same underlying OM subunit (Fig. 5f). However, any possible interactions between prefusion G and OM during assembly would disappear when the G trimer transitions to the postfusion conformation, as indicated by the location of the super-complex (i.e., off the OM) (Fig. 5f) and the non-specificity of the orientation between the postfusion G trimer and the underlying OM lattice (Extended Data Fig. 4).

## Discussion

In this study, we have used helical cryoEM sub-particle reconstruction and cryoET subtomogram averaging to determine the first near-atomic resolution *in situ* structures of M and N of VSV, as well as the distribution of G trimers on the VSV membrane envelope. These *in situ* structures establish the mode of molecular interactions among genomic RNA, N, M and G. Aside from the significance embodied in these long sought-after structures are two transformational discoveries. First, the stoichiometry between N and M is not the previously thought 1:1, but 1:2; a double-layer of M, composed of IM and OM, surrounds a layer of N. Second, the G trimers on the viral membrane envelope are organized in pseudo-crystalline lattices, with potential tenuous interactions with OM through their endodomains.

NSVs all contain structural proteins N and M. A careful comparison of subunit interactions in the nucleocapsid of VSV with those of MeV indicates that their respective shapes—the characteristic bullet shape of VSV and the helical shape of MeV—are influenced by N-N interactions (Extended Data Fig. 5). In MeV, the assembly of the helical nucleocapsid relies entirely on N-N interactions, both vertically and laterally^24^. Each N subunit interacts directly with an adjacent N by an α helix of the N-arm (Extended Data Fig. 5, right), which is the configuration that gives rise to its slimmer helical nucleocapsid. In VSV, however, the flexible β-strand hairpin of the N-arm from N_U-1_ anchors on the C-loop of N_U+1_ (Fig. 3b); the flexibility of the involved N-arm and C-loop permits the different curvatures of N helical turns from the tip to the trunk (Fig. 3 a, b). This flexibility also explains how different numbers of asymmetric units per helical turn are present in different VSV virions. In addition to the interactions between N subunits, other interactions, including RNA-N, N-IM, and IM-OM, make the large (∼700 Å diameter), three-layered nucleocapsid sufficiently stable for packaging numerous polymerase complexes. Notably, the vertical interactions between N subunits in VSV are minimal: only IM subunits from successive turns interact with each other through a flexible N-terminal segment. The IM helix directly interacts with the N helix, with each IM positioned between two helical turns of N and stabilizing the overall structure.

One of the primary functions of M is to drive virus budding. A network of M molecules is associated with the inner leaflet of the virus envelope and the endodomains of viral glycoproteins, as shown for Newcastle disease virus (NDV)^25^, MeV^26^, Ebola/Marburg viruses^27^, and influenza viruses^28^. However, the other primary function of M, to stabilize the nucleocapsid, is not fully understood. In the structure reported here, VSV is shown to have a double layer of M instead of a single layer, as previously thought^20,29^. Compared to the IM subunit, the OM subunit has a 40° rotation along the radial axis and an additional 40° rotation along the vertical axis (Extended Data Fig. 6b). Although IM and OM have similar structures, this large rotation of OM makes the outer-facing surfaces of IM and OM quite different (Extended Data Fig. 6 c, d). Generally, the outside surface of IM is more hydrophilic, while that of OM is more hydrophobic (Extended Data Fig. 6 c, d). In addition, the rotation of OM changes the interactions that are present in the IM layer (Fig. 2i and Extended Data Fig. 1b), which allows OM subunits to be distributed differently from IM. In the cryoET density, we noticed that each OM subunit has a small region attached to the membrane envelope (Extended Data Fig. 6a, red arrows); further analysis indicated that the attachment region involves two protruding loops, Pro195 – Ser199 and Ala216 – Gly218 (Extended Data Fig. 6d, green). Viewed from the side, these two loops are in a close position to interact with the membrane. In both structures of the prefusion and postfusion G trimers^18,19^, 80 amino acids of G at the C-terminal, which constitute the membrane-proximal region, the transmembrane domain and the endodomain, are removed by protease digestion. The endodomain of G contains as many as seven positively charged residues (Lys and Arg) (Extended Data Fig. 6e). Interestingly, surface potential analysis of OM shows that a negatively charged region is located in the middle of the relatively hydrophobic surface of OM (Extended Data Fig. 6d). The negatively charged region comprises a small helix Val189 – Tyr192, in which the side chain of Glu191 and the main chain contribute to the negative potential. Thus, the endodomain of G could reasonably interact with this region of OM.

The double layer of M may be the result of its dual role in virion assembly; the functional explanation for the double layer of M is supported by other studies. The OM layer in VSV is recognized to associate with the viral membrane and the endodomain of the glycoproteins, as M does in other viruses^26,30^. Membrane-derived M proteins (presumably OM) have been shown to bind nucleocapsids condensed by M proteins (presumably IM), but not intracellular nucleocapsids, which only contain RNA and N^31,32^. This binding (or lack thereof) may be explained by the different orientations, and thus the different binding affinities, of OM and IM to N. Cytosolic M protein can bind with intracellular nucleocapsids, but with much lower affinity than membrane-derived M proteins to M-condensed nucleocapsids. Without association with the M protein, the nucleocapsid was in an extended state^33,34^. The assembly of a bullet-shaped virion requires functional M proteins^35^, presumably IM subunits for condensing the nucleocapsid. In other NSVs, the nucleocapsid self-assembles into a condensed helical structure ready for packaging in a virion. Their M proteins also bind the nucleocapsid prior to its incorporation into the virion^30,36-38^. Since formation of the helical nucleocapsid in other NSVs does not require association with the M protein, it is not clear if a secondary M subunit participates, in either a partial or full layer, in bridging the membrane-bound M layer and the packaged nucleocapsid in those virions.

The extensive interactions between N-IM and between IM-OM revealed by our *in situ* structures, as well as the interactions between OM and G proposed by our cryoET results, suggest a mechanism of VSV assembly as follows (Fig. 6a). After genome replication, the nucleocapsid tends to assemble into a bullet-shaped structure^35,39^, assisted by the association of IM. IM binding prohibits the nucleocapsid from serving as the template of viral synthesis and allows it to be transported to the plasma membrane. Meanwhile, M molecules, capable of associating with the host cell membrane^40^, form a regular mesh on the membrane that enables binding with the IM-stabilized nucleocapsid. Membrane microdomains containing G clusters that presumably share a lattice similar to that in the crystal of prefusion G trimers^19^ have been observed, and the IM-decorated nucleocapsid would bind underneath such microdomains through IM-OM interactions. This association would introduce curvature to the microdomain, leading to budding of an infectious virion. The pseudo-crystalline organization of G trimers with predominant hexagonal and occasional pentagonal tiles suggests that there are interactions between OM and the endodomain of G. Therefore, the atomic details unveiled by these *in situ* structures account for not only the genesis of the bullet-shaped nucleocapsid but also the recruitment of meta-stable trimers of G to form a complete infectious virion. Finally, based on current structural and biological information, we constructed a pseudo-atomic model of an entire VSV virion, which contains 1235 N, 1235 IM, 1235 OM, 11115 RNA nucleotides, 49 LP_2_, and numerous G-trimers (Fig. 6b-e and Supplementary Video 4, 5).

**Fig. 6.**
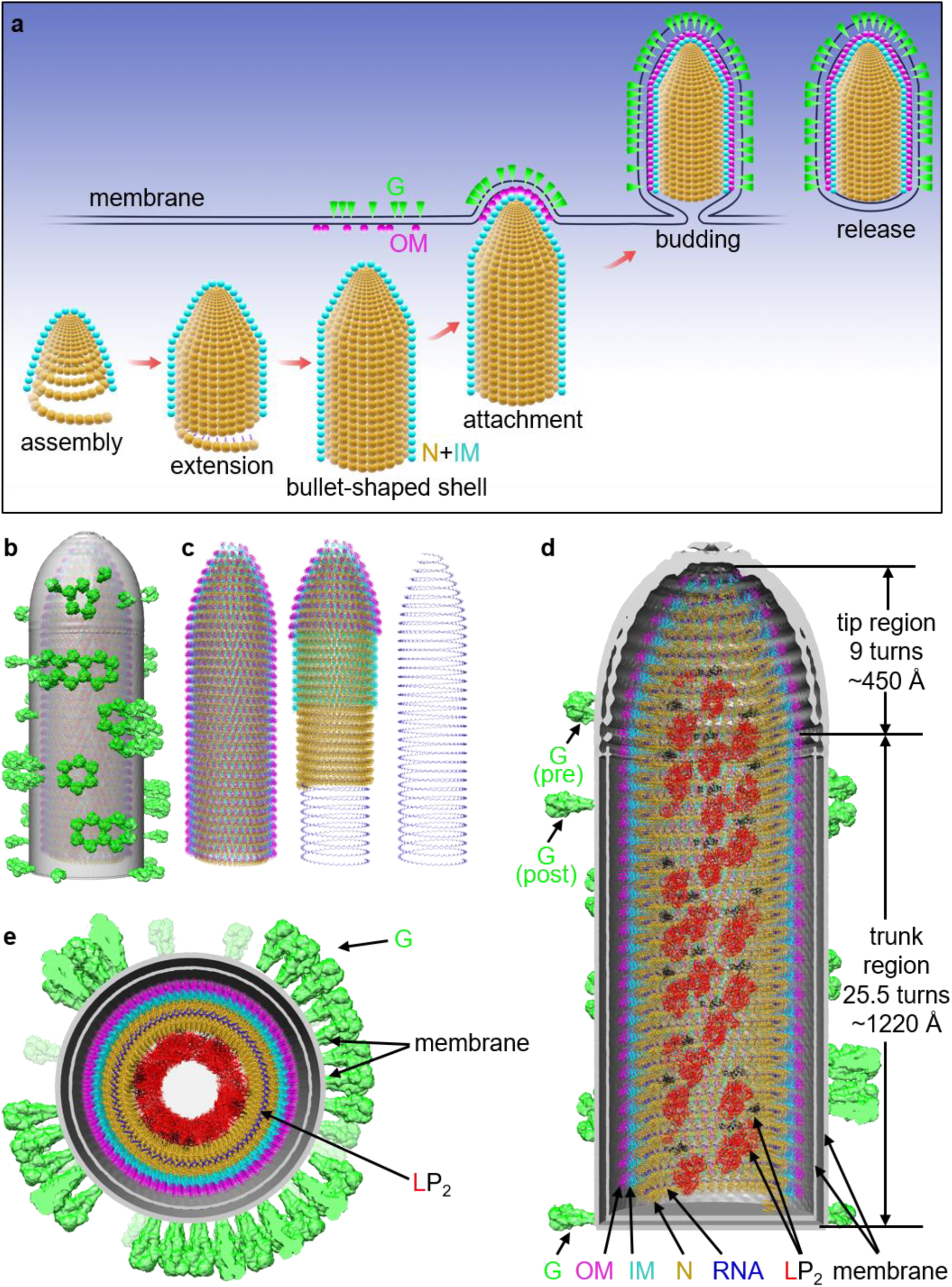
VSV assembly model and reconstruction of an entire VSV virion. **a**, In the cytosol, the nucleocapsid, stabilized by IM binding, is wound into a conical tip at the 3’ end. As assembly continues, the bullet-shaped shell of the nucleocapsid is formed, and more IM is bound. Meanwhile, OM is attached to the plasma membrane and colocalized with membrane-anchored G trimers. At the budding stage, the tip of the bullet-shaped shell attaches to the OM on the membrane, which induces curvature of the membrane and initiates budding. **b**, Overall view of the entire VSV model, which includes 1235 N, 1235 IM, 1235 OM, 11115 RNA nucleotides, 49 LP_2_, and numerous G-trimers. **c**, Overall view of the M-coated capsid. Left: all proteins in the M coated capsid shown; middle: layers are hidden by segment; right: RNA shown alone. **d**, Cross section of (**b**) viewed from side; each component is labeled. **e**, Cross section of (**b**) viewed from tip.

From a technical point of view, our work highlights an innovative approach of sub-particle reconstruction for filamentous objects with flexible helical parameters. The workflow complements recent successes of focused refinement of RELION^41^ to improve resolution of flexible local regions of a complex and sub-particle reconstruction to resolve asymmetric structures in icosahedral viruses^42,43^. This method should be generally applicable to other helical/filamentous assemblies that are common in viruses and cells.

## Data Availability

The sub-particle reconstruction and cryoET averaged density maps are deposited to the Electron Microscopy Data Bank under the accession numbers of EMD-XXXXX and EMD-XXXXX. The atomic model is deposited in the Protein Data Bank under the accession numbers of XXXX, and Electron Microscopy Data Bank under the accession numbers of EMD-XXXXX, respectively.

## Acknowledgements

We thank Titania Nguyen for editing the manuscript. This project was supported in part by grants from the US NIH (GM071940 and AI094386 to Z.H.Z.) and grant from University of Science and Technology of China (the Fundamental Research Funds for the Central Universities WK9100000025 to K.Z.). We acknowledge the use of resources at the Electron Imaging Center for Nanomachines supported by UCLA and grants from the NIH (1S10OD018111 and 1U24GM116792) and the National Science Foundation (DBI-1338135 and DMR-1548924).

## Author contributions

Z.H.Z., M.L. and P.G. conceived the project; J.T. cultured VSV with supervision of M.L.; K.Z. purified VSV and prepared cryoEM and cryoET grids; K.Z. recorded cryoEM images; K.Z. and Z.S. recorded cryoET tilt series; P.G. developed the helical sub-particle reconstruction method and worked together with K.Z. to process the cryoEM data; K.Z. built atomic models; Z.S. performed cryoET reconstructions and subtomogram averaging; Z.H.Z., K.Z., Z.S. and M.L. interpreted results and wrote the paper; all authors edited and approved the paper.

