## Extended Data for "Atomic model of Vesicular Stomatitis Virus and Mechanism of Assembly"

### Supplementary Information

#### Materials and Methods

##### Isolation of VSV virions

VSV virions were cultured as previously described[1], and full VSV particles were isolated from media by density-gradient centrifugation. The virions were pelleted at 30,000g for 2 hours and resuspended in phosphate buffered saline (PBS, pH 7.4). The stock was then subjected to another low-speed centrifugation at 12,000g for 5 minutes to remove large aggregates. The resulting suspension was loaded on a 12ml density gradient containing 0-50% potassium tartrate and 30-0% glycerol (prepared by 6ml 50% potassium tartrate and 6ml 30% glycerol). After centrifugation at 40,000g for 1 hour, the VSV-containing band was extracted with a syringe and slowly diluted in 10 ml PBS. The dilution was pelleted at 30,000g for 2 hours; the subsequent pellet was kept on wet ice for 4 hours and resuspended in a small volume of PBS prior to use for cryoEM and cryoET sample preparation.

**CryoEM sample preparation, movies acquisition and drift correction**

For single-particle cryoEM, each aliquot of 2.5 μl of the purified sample was applied onto a glow-discharged holey copper grid (300 mesh, Quantifoil R 1.2/1.3, Ted Pella). The grid was blotted and flash-frozen in liquid ethane with an FEI Mark IV Vitrobot. An FEI TF20 cryo-EM instrument was used to screen grids and optimize the conditions.

Optimized cryo-EM grids were loaded into a Titan Krios 300 kV electron microscope (Thermo Fisher Scientific) equipped with a Gatan imaging filter (GIF) Quantum LS and a Gatan K2 Summit direct electron detector. Movies were acquired with Leginon[2] in super-resolution mode at a nominal magnification of 105,000x (yielding a calibrated pixel size of 0.66 Å at the specimen level). The GIF slit width was set to 20 eV. A total number of 45 frames were acquired in 9 seconds for each movie, giving a total dose of ~60 $e^-/Å^2$/movie.

Frames in each movie were aligned for drift correction with the graphics processing unit (GPU)-accelerated program MotionCor2 (ref. [3]). The first and last frame were discarded during drift correction. Two averaged micrographs, one with dose weighting and the other one without dose weighting, were generated for each movie after drift correction. The averaged micrographs were binned 2x2 to yield a pixel size of 1.325 Å. The micrographs without dose weighting were used for CTF estimation and particle picking, while those with dose weighting were used for particle extraction and in-depth processing.

**2D and 3D classification**

The data processing workflow for VSV cryoEM dataset is summarized in Extended Data Fig. 7. The CTF estimation of each micrograph was performed by CTFFIND4 (ref. [4]). From a total of 2,008 micrographs, 1,692 good ones were selected and phase-flipped by Bsoft[5]. 7,732 VSV virions (start-end coordinates pair) were then picked up manually using RELION[6].

With RELION's helical extraction function, each virion was sub-divided into overlapping segments with dimensions of 800 × 800 square pixels and an inter-box distance of 100 Å. 97,921 particles were extracted and 2× binned to 400 × 400 square pixels (pixel size: 2.65 Å) to speed up further data processing. These particles were then subjected to reference-free 2D classifications. Classes with bad particles (i.e., classes with fuzzy or uninterpretable features) and those containing either the virion tip region or the bottom region were discarded, yielding a total of 72,923 selected particles.

To account for the variable numbers of asymmetric units per helical turn, we sorted particles according to their helical parameters through four rounds of 3D classification. The first three rounds generated the density maps used as the initial references, with which we subsequently sorted the particles into different classes in the fourth and final round through multi-reference 3D classification. In the first round of 3D classification, we used *relion_helix_toolbox* with known and calculated helical parameters of VSV and the atomic models of N[7] (PDB: 2WYY) and M[8] (PDB: 2W2R) to generate 6 reference models. These models ranged from 35.5 to 40.5 asymmetric units per turn with an interval of 1. To calculate the helical parameters (helical rise and helical turn) of these models, we consulted a previous study[7] to find the distance between two adjacent turns (helical pitch), which is about 50 Å. Thus, the helical rise is 50 Å / units per turn, and the helical turn is -360° / units per turn (a negative degree indicates left-handedness of the helix). Each segment was transferred to a density map (20 Å low-pass filtered) using the *molmap* command in Chimera[9], and the 6 resulting density maps were used as the initial references for 6 independent 3D classification jobs. In each job, all good particles were employed and helical parameters imposed according to the number of asymmetric units per turn. 3 classes were asked for each job, and no helical parameter search was applied. The reconstructed map with the best features among the 3 classes was selected as the reference for the second round of classification; this was repeated for each job. In total, 6 maps were selected from 6 independent 3D classifications. In the second round of 3D classification, a single Star file was used, containing the 6 maps selected from the first round and their corresponding helical parameters as the initial references. All good particles were employed again, resulting in 6 classes and 6 corresponding maps. In the third round, each class, containing a particle subset and map, from the second round was selected and subjected to an independent 3D classification job. For each job, 3 classes were asked and helical parameters were searched for within a very small range. Thus, the particles were further classified, leading to classes with improved reconstructed maps and more accurate helical parameters. However, for the job related to 40.5 asymmetric units per turn, we noticed that one of the 3 classes had much lower resolution compared to the other two, suggesting the existence of particles with 41.5 asymmetric units per turn. Thus, we performed additional 3D classification jobs similar to those in the second round for this particle subset, resulting in a reconstructed map of and refined helical parameters for particles with 41.5 asymmetric units per turn. Through the three rounds of classifications above, 7 good initial reference maps and their corresponding helical parameters were prepared. In the fourth round, these improved maps and helical parameters were used as references for a multi-reference 3D classification job. All good particles were employed, and 7 classes with variable numbers of asymmetric units per turn (35.5-41.5) were identified, with particle distribution percentages of 6.8%, 15.1%, 12.6%, 31.8%, 20.7%, 6.0% and 7.0%, respectively (Extended Data Fig. 7 and Supplementary Table 2).

**Sub-particle reconstruction**

3D classification resulted in 7 classes with different helical parameters; however, further 3D auto-refinement of each class failed to achieve high resolution, which indicated local flexibility inside the helix. Subsequent data processing focused on the major class, which has 38.5 asymmetric units per turn and comprises 31.8% of the total number of particles. To accommodate the helix's flexibility and improve resolution, we re-extracted small patches from the helical segments as sub-particles (Extended Data Fig. 7) with a custom-designed program called *helisub.C*. The location of each sub-particle was calculated using the refined helical parameters (helical rise: 1.30 Å, helical turn: -9.35°) and the measured radius (210 Å) of the helical trunk. To avoid duplicates, 60 sub-particles from each segment's central helical turns were extracted. In total, 1,392,420 sub-particles were extracted from 23,207 maternal segments, with a box size of 160x160 (unbinned). Along with the image stack of all the sub-particles, a new data Star file, containing the Euler angles and offsets of each sub-particle, was also generated. The initial reference was directly obtained from the image stack of the sub-particles utilizing the *relion_reconstruct* command; the sub-particles were then subjected to a 3D auto-refinement (local search), yielding a density map at 3.47 Å resolution (Extended Data Fig. 8).

The effective resolution was estimated based on the ''gold standard'' refinement procedures and the 0.143 Fourier shell correlation (FSC) criterion (Extended Data Fig. 8b). Local resolution was estimated using Resmap[10] (Extended Data Fig. 8c).

**Atomic modeling and model refinement**

Atomic model building was accomplished in an iterative process involving Chimera[9], Coot[11] and Phenix[12]. First, the crystal structures of VSV matrix Protein M[8] (PDB: 2W2R) and nucleoprotein N[13] (PDB: 2GIC) were fitted into the cryoEM map by Chimera. Fitting revealed an extra density corresponding to single stranded RNA, whose model was subsequently built manually with Coot. This initial model was then refined using Phenix in real space with secondary structure and geometry restraints. Subsequently, the resulting model was manually adjusted for with Coot by fixing those residues not matching the cryoEM density. The Phenix and manual refinements were repeated until no improvement was possible. Refinement statistics of the VSV nucleocapsid are summarized in Supplementary Table 3. The model was also evaluated based on MolProbity scores[14] and Ramachandran plots (Supplementary Table 3).

**CryoET sample preparation, movies acquisition and drift correction**

For cryoET, fiducial gold beads of 5nm size were added into the purified VSV sample at a ratio of 1:20 (V/V). Each aliquot of 2.5 μl of the sample was applied onto a glow-discharged holey copper grid (200 mesh, Quantifoil R 3.5/1, Ted Pella). The grid was blotted and flash-frozen in liquid ethane with an FEI Mark IV Vitrobot. An FEI TF20 cryo-EM instrument was used to screen grids and optimize the conditions.

Optimized cryo-EM grids were loaded into the same Titan Krios 300 kV electron microscope (Thermo Fisher Scientific) equipped with a Gatan imaging filter (GIF) Quantum LS and a Gatan K2 Summit direct electron detector. The tomography data collection was performed with SerialEM[15] software v3.8, with a dose-symmetric scheme[16]. The beam was aligned in nano-probe mode and the GIF slit width was set to 20 eV. Tilt series movies were recorded in dose-fraction mode, with a nominal magnification of 81,000X, calibrated pixel size of 1.74 Å, counting mode, defocus range 2.5~3.5um, tilt range -60°~60°, tilt increment 3° with constant dose, total dose ~120 $e^-/Å^2$. 30 tilt series were acquired in total. Frames (typical 5~7 frames per movie, 0.2 seconds per frame) in each movie were then aligned for drift correction with the graphics processing unit (GPU)-accelerated program MotionCor2 (ref. [3]). The averaged micrographs were used for the following tomogram reconstruction.

**Tomogram reconstruction and 3D denoising**

Fiducial-based tilt series alignment and tomographic reconstruction were performed with IMOD[17] v4.9 software package. Gold beads were manually picked and automatically tracked. For each tilt series, aligned tilt images were 4 times binned and 2 times binned for tomogram reconstruction by weighted back projection, respectively. The 4 times binned tomograms were reconstructed for particle picking, and the 2 times binned tomograms for subtomographic averaging. The defocus value of each tilt images was estimated by CTFFIND4 (ref. [4]).

For better visualization, 5 times binned tomograms were also reconstructed for denoising by Noise2map in Warp[18]. For each tilt series, even-numbered and odd-numbered tilt images were separated and reconstructed by weighted back projection, respectively. The pair of tomograms were trained about 10000 iterations for a noise model, which was then used to generate a denoised tomogram. Additionally, 5 times binned tomograms with SIRT-like filter, equivalent to those reconstructed by SIRT algorithm with 10 iterations, were also reconstructed for comparison and visual inspections.

**Virus classification based on numbers of asymmetric units per helical turn**

Due to the heterogeneity of VSV particles, virions were grouped into 7 classes according to their number of asymmetric units per turn. In total, we manually selected 124 virions, and each of them was registered by a vector pointing from the blunt end to the tapered end. To simplify subsequent processing, only their trunk regions were taken into consideration in the following. The helical parameter solved from the previous cryoEM helical reconstruction consists of 37.5 asymmetric units per turn and a 50.8 Å rise along the helical axis[7]. The seven sets of helical parameters we used for virus sorting have identical helical rise, but the number of asymmetric units per turn ranges from 35.5 to 41.5. For each trunk, seven left-handed helical samplings, each with a different units-per-turn measurement, were applied to generated initial positions and orientations on the cylindrical lattice, which were then used to extract subtomograms. With these defined initial orientations, seven reconstructions were obtained after averaging without search. Among the seven reconstructions, only the one which coincided with the helical parameter set of each individual virion showed the strongest features, while others were smeared due to mismatching. The class which had 38.5 asymmetric units per turn comprised the majority, with 57 out of 124 virions belonging to this class. Thus, virions with 38.5 asymmetric units per turn were considered in the following particle picking and subtomogram averaging.

**Particle picking (G protein) for subtomogram averaging**

The averaged structure of G proteins in postfusion conformation[19] was low-pass filtered to 40 Å and used as initial template. 4x binned tomograms were cropped into sub-regions, with each sub-region containing only one virion. Template matching was performed with *templateSearch* function in emClarity[20]. For each sub-region, 400 highest-scoring cross-correlation peaks were extracted, with a minimal distance of 54Å (radius of G in postfusion conformation) between peaks to prevent multiple detections. To remove obvious false positives, such as the fiducial gold beads and the M proteins inside the virions, geometric constraints based on the bullet (cylindrical) shape of the virions were applied to exclude the peaks which were not on the viral membrane. 9443 subtomograms from 57 virions were retained and then refined in Relion[21].

We then extracted subtomograms with a box size of 180 from 2 times binned tomograms with the coordinates of subtomograms derived from template matching. The orientation of each subtomograms has three Euler angles calculated from the reported orientations of template match and denoted as parameters within the Relion Star file: rot (-rlnAngleRot), tilt (-rlnAngleTilt) and psi (-rlnAnglePsi). With the pre-determined orientations, an initial reference was obtained by *relion_reconstruct*. The refinement converged into a density map of G in postfusion conformation of ~15 Å. In order to further remove false positives, especially the subtomograms containing only the membrane but no G, a skip-align classification was performed with a tight mask round the ectodomain of G. Two classes were claimed here, and the class without the ectodomain of G was excluded. Eventually, 6030 subtomograms from 57 virions, which have 38.5 asymmetric units per helical turn, were retained and used in the following analysis.

**Subtomographic averaging and 3D classification**

As for the subtomograms averaging of G, local searches from auto-sampling of 1.8 degrees and 4 pixels offset searching range were performed by Relion[6], with three-fold symmetry imposed during the refinement. The final resolution was estimated with two independently refined maps from two halves of the dataset by *relion_postprocess* and determined to be 14.9 Å at 0.143 criteria with gold standard FSC.

For the averaging of G proteins with different numbers of neighbors, the workflow is summarized in Extended Data Fig. 2. A new Star file was generated after C3 symmetry was released by *relion_particle_symmetry_expand* and 3 times of numbers of subtomograms were subject to a skip-aligned 3D classification, with a mask around one of the three neighbors (Extended Data Fig. 2 a, b). Two classes were claimed here. Based on how many symmetrized copies belonged to the first class (with neighboring G protein), all original subtomograms were divided into 4 groups: subtomograms whose 0, 1, 2, 3 symmetrized copies went to the first class originally had 0, 1, 2, 3 neighbors adjacent to the central G protein, respectively (Extended Data Fig. 2 a, b). 4 groups of G with 0, 1, 2 and 3 neighbors contained 918 (15.2%), 2581 (42.8%), 2004 (33.2%) and 527 (8.8%) subtomograms, respectively. The groups were then refined individually, with C3 symmetry imposed for groups of 0 and 3 neighbors and C1 for groups of 1 and 2 neighbors. The final resolutions of these 4 averaged maps were also estimated by *relion_postprocess* and determined to be 18.4 Å, 19.6 Å, 20.9 Å and 20.9 Å at 0.143 criteria with gold-standard FSC (Extended Data Fig. 2c).

**Reconstruction of an entire bullet-shaped virion structure**

We divided the nucleocapsid of the virion into tip and trunk regions and reconstructed the trunk region of the virion first. The trunk region is a regular helix, with the same diameter and helical parameters per turn. Based on the asymmetric unit structure (1N, 1 IM, 1OM and 9 nucleotides) described above and the known helical parameters of a 38.5 subunit/turn helix (helical twist = -9.35°, helical rise = 1.30 Å), a reconstructed model of 120 asymmetric units was constructed by RELION[6] (*relion_helix_toolbox*). The model fit perfectly into the cryo EM density of the major class from helical sub-particle reconstruction (class 4, 38.5 subunits/turn) (Extended Data Fig. 9a).

Next, we reconstructed the tip region. To find the number of subunits per turn and the subunits' relative orientation in the tip, we manually picked 9,515 tips from all micrographs and carried out a 2D classification (Extended Data Fig. 9b). Based on previous studies[7] and the 2D classification data, we concluded that the tip region is composed of a spiral whose diameters increase turn by turn from the pole. In addition, the angles between the long axis of the asymmetric unit in the spiral and the axis of the trunk increase gradually as the spiral proceeds from the pole (Extended Data Fig. 9c). Remarkably, we noticed that the distance between turns in the tip region remains the same (~50 Å). To simplify the geometric model of the tip region, we treated each turn as a standard helical turn.

As we already knew that each asymmetric unit corresponds to 9 RNA nucleotides and the RNA strand is uninterrupted, we used the RNA strand as a good reference to calculate the diameter of each turn in the tip and subsequently the number of asymmetric units. We projected the major class from 3D classification of the trunk (including two membrane layers) and the aforementioned trunk model (Extended Data Fig. 9c) to get the side view of the trunk. By aligning the projection images of the trunk and the 2D classification of the tip, we were able to locate the position of the RNA strand inside the 2D classification of the tip (Extended Data Fig. 9c, yellow dashed lines). Subsequently, we sketched the membrane curve and defined the edges of the RNA strands by translating the curves to both sides (Extended Data Fig. 10a). We then measured the diameters of the RNA and calculated the subunit numbers of each turn (Extended Data Fig. 10a and Supplementary Table 4). Further analysis showed that the diameter of each turn increases from the pole, while the size of that increase decreases. With the distance between adjacent turns and the difference in diameter between adjacent turns, we could calculate the slope of tangent line, which indicates the tilt angle of subunits in each turn except for the first turn (Extended Data Fig. 10b, and Supplementary Table 4). Thus, we were able to reconstruct each turn of the tip region. Using these turns, as well as manual refinement, we reconstructed the full tip region, which includes 9 turns. Together with the first turn of the trunk, these 10 turns contain 291 subunits (Extended Data Fig. 10c). Eventually, we joined the trunk region and tip region together as a full length nucleocapsid model containing 1,235 subunits; each subunit contains 1 N, 1 IM, 1 OM and 9 RNA nucleotides (Fig. 6c).

Placement of $LP_2$ inside the nucleocapsid and G trimers on the envelope are based on the cryoET reconstruction in this study and previous study[22], resulting in a pseudo-atomic model of the entire virion (Fig. 6b, d, e).  The PDB file and associated MRC density maps, together with a Chimera[9] session (.py file) to display them, are provided as a supplementary data file. The program for data processing is provided in Github.

**Graphics Visualization**

Visualization of the density maps, atomic models, figures and movies, are made by UCSF Chimera[9] and IMOD[17].

#### Extended Data Figures:

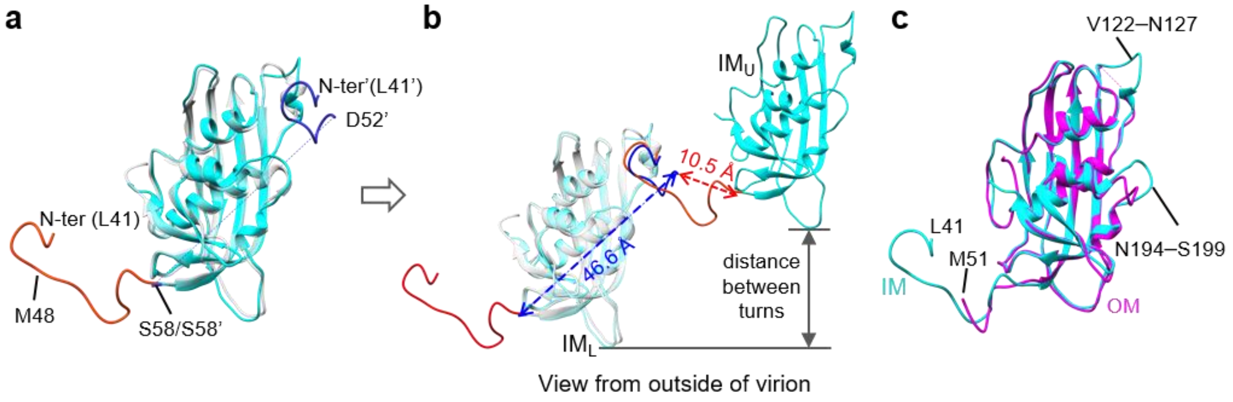


**Extended Data Fig. 1 | Structure comparison of M. a:** Superposition of the *in situ* structure of M (IM) and crystal structure of M. The *in situ* IM is colored in cyan except for the N-termini, in red, while the M in crystal structure is colored in transparent gray except for the N-termini, in blue. **b**: Distance between terminal residues of IM in cryo EM structure and the determination of N-termini affiliation. 7 residues (D52-S58) were missing in the structure. Between the blue N-terminal segment and the $IM_U$ is a distance of 10.5 Å, which can be bridged by those 7 residues; between the same segment and the $IM_L$ is a distance of 46.6 Å, which cannot. Therefore, the blue N-terminal segment must belong to $IM_U$. **c:** Superposition of the inner M (IM) and outer M (OM) subunits. IM is colored in cyan and OM in magenta.

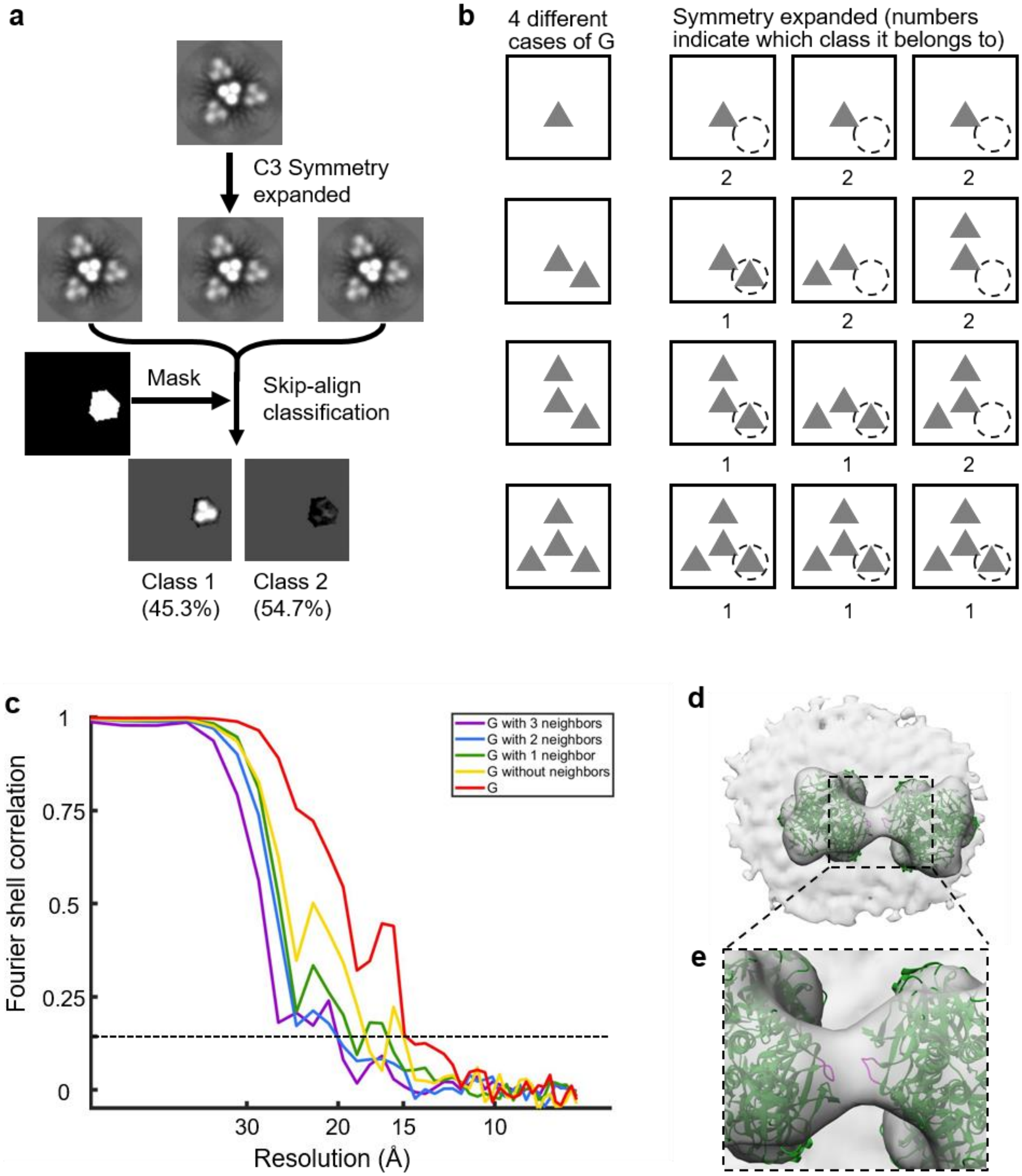


**Extended Data Fig. 2 | CryoET analysis for VSV G. a, b:** Workflow of 3D classification of G trimers with different numbers of neighboring G trimers. **c:** Fourier shell correlation coefficient as a function of spatial frequency of subtomogram averages of G with different numbers of neighboring G trimers. The dashed line depicts the 0.143 resolution cut-off. Red: average of G regardless of number of neighboring G trimers, 14.9 Å; yellow: average of G without neighboring G trimers, 18.4 Å; green: average of G with one neighboring G trimer, 19.6 Å; blue: average of G with two neighboring G trimers, 20.9 Å; purple: average of G with all three neighboring G trimers, 20.9 Å. **d:** Subtomogram averaged structure of G with one neighboring G trimer fitted with atomic models of the crystal structure of G in postfusion conformation, showing the interactions between two G trimers. **e:** Zoom-in view of the box region in (**d**). N-termini of G (residues 8-14), colored magenta, are potentially involved in the interaction.

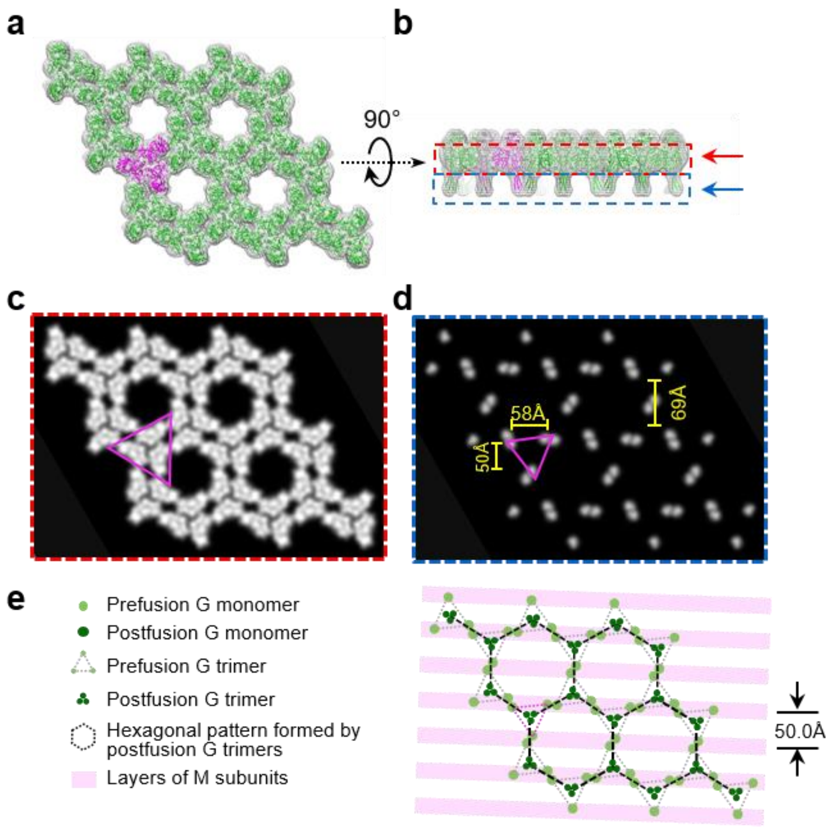


**Extended Data Fig. 3 | 2D hexagonal lattice in crystals of prefusion G. a, b**: Density maps derived from atomic models of prefusion G trimers showing the 2D hexagonal lattice in crystals. A density map is filtered to 15 Å. Models colored as magenta are one of the prefusion trimers. (**a**) is the top view; (**b**) is the side view. **c, d:** top views of the density map from different sections, indicated as arrows in (**b**) (red arrow for **c**, blue arrow for **d**). The magenta triangle indicates the same prefusion trimer in (**a**) and (**b**). Distances between the fusion loops of two G protomers, the equilateral triangle height of the G trimer and two centers of G trimers, are measured as 58 Å, 50 Å and 69 Å, respectively. **e:** Illustration of the 2D lattice formed by G trimers on the viral membrane. The organization of G matches with that of OM inside the membrane.

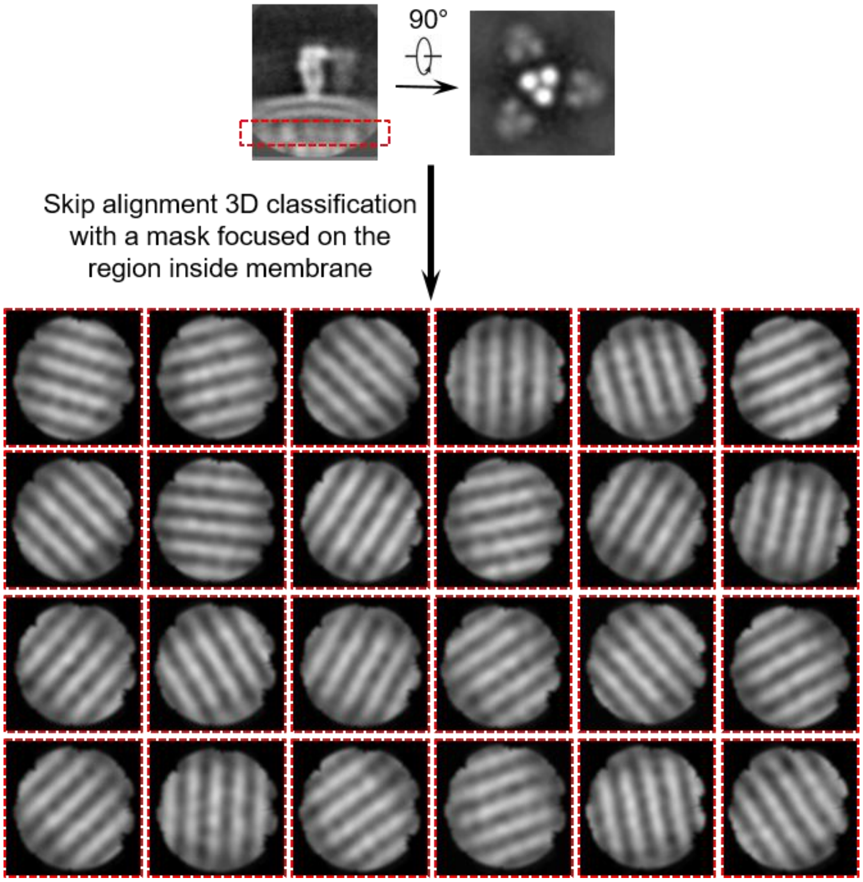


**Extended Data Fig. 4 | 3D classification showing different orientations of RNP inside the viral membrane with respect to the postfusion G trimers outside the viral membrane.**

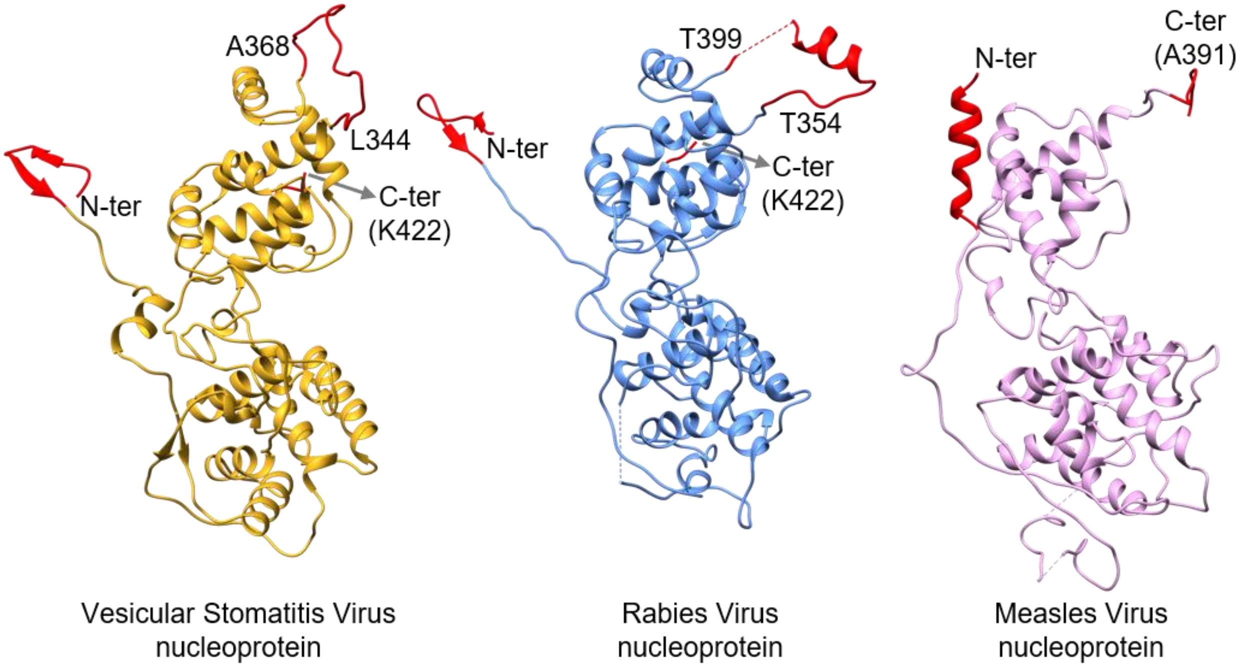


**Extended Data Fig. 5 | Structure comparison of nucleoproteins (N) from VSV, rabies virus and measles virus.** The N-termini, C-termini and C-loop are highlighted in red.

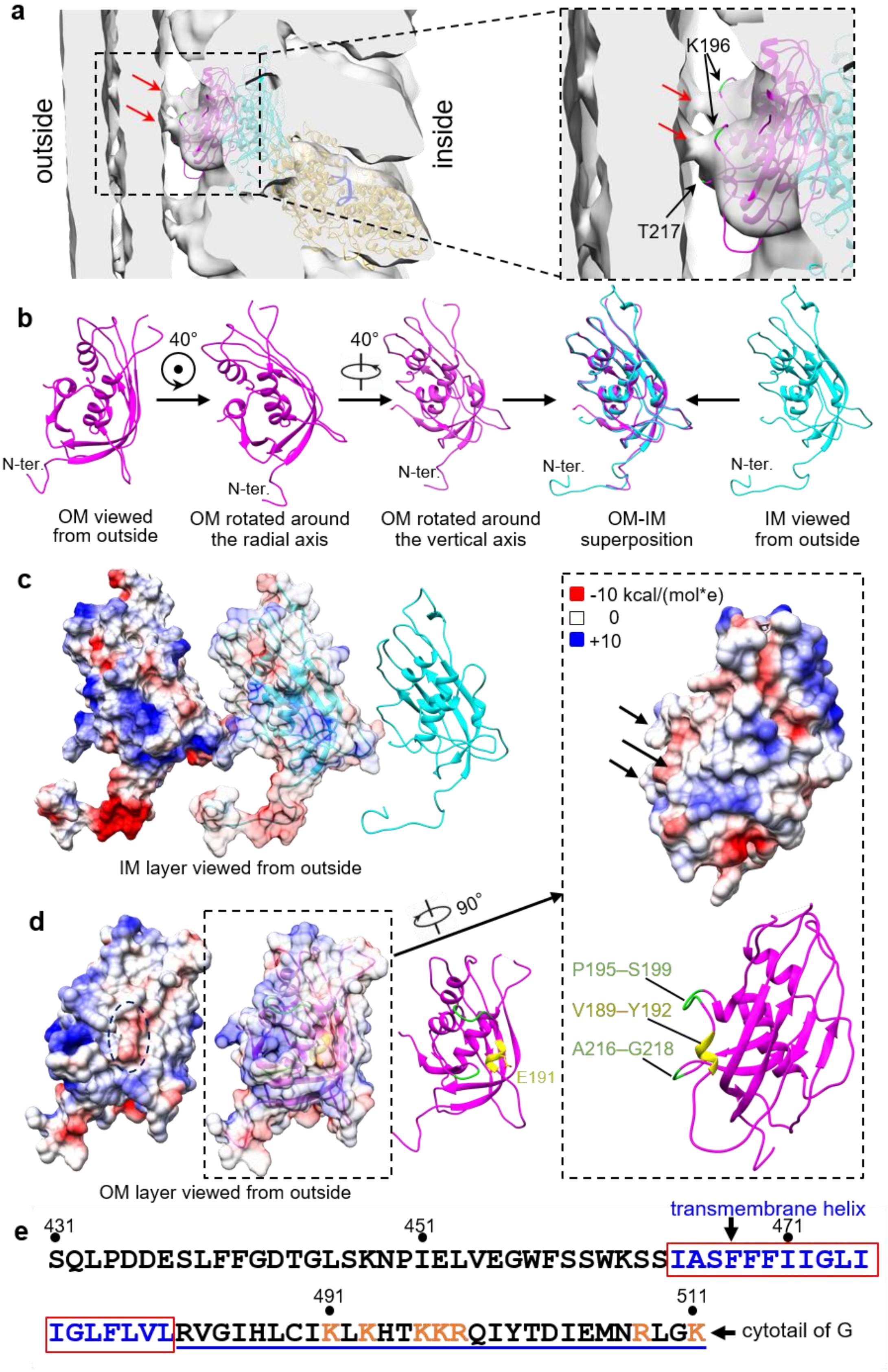
a
outside
inside
K196
T217
b
40°
40°
N-ter.
N-ter.
N-ter.
N-ter.
N-ter.
OM viewed from outside
OM rotated around the radial axis
OM rotated around the vertical axis
OM-IM superposition
IM viewed from outside
c
-10 kcal/(mol*e)
0
+10
IM layer viewed from outside
d
90°
E191
P195–S199
V189–Y192
A216–G218
OM layer viewed from outside
e
transmembrane helix
431
451
471
SQLPDDESLFFGDTGLSKNPIELVEGWFSSWKSSIASFFFIIGLI
491
511
IGLFLVLRVGIHLCIKLKHTKKRQIYTDIEMNRLGK
cytotail of G

**Extended Data Fig. 6 | Interactions between G trimers and between OM and viral membrane. a:** Subtomogram averaged structure of the nucleocapsid fitted with the atomic models solved above, showing the thin densities linking the OM and the inner leaflet of the viral membrane. Red arrows indicate the linking densities. **b**: Step-by-step superposition between OM and IM subunits. **c**: Outside surface potential of IM. **d**: Outside surface potential of OM. The negatively charged region is marked by dashed lines; a small helix corresponding to this region is colored in yellow.  Two loops located close to the membrane are colored in green. **e**: The amino acid sequences of 80 C-terminal residues of G.

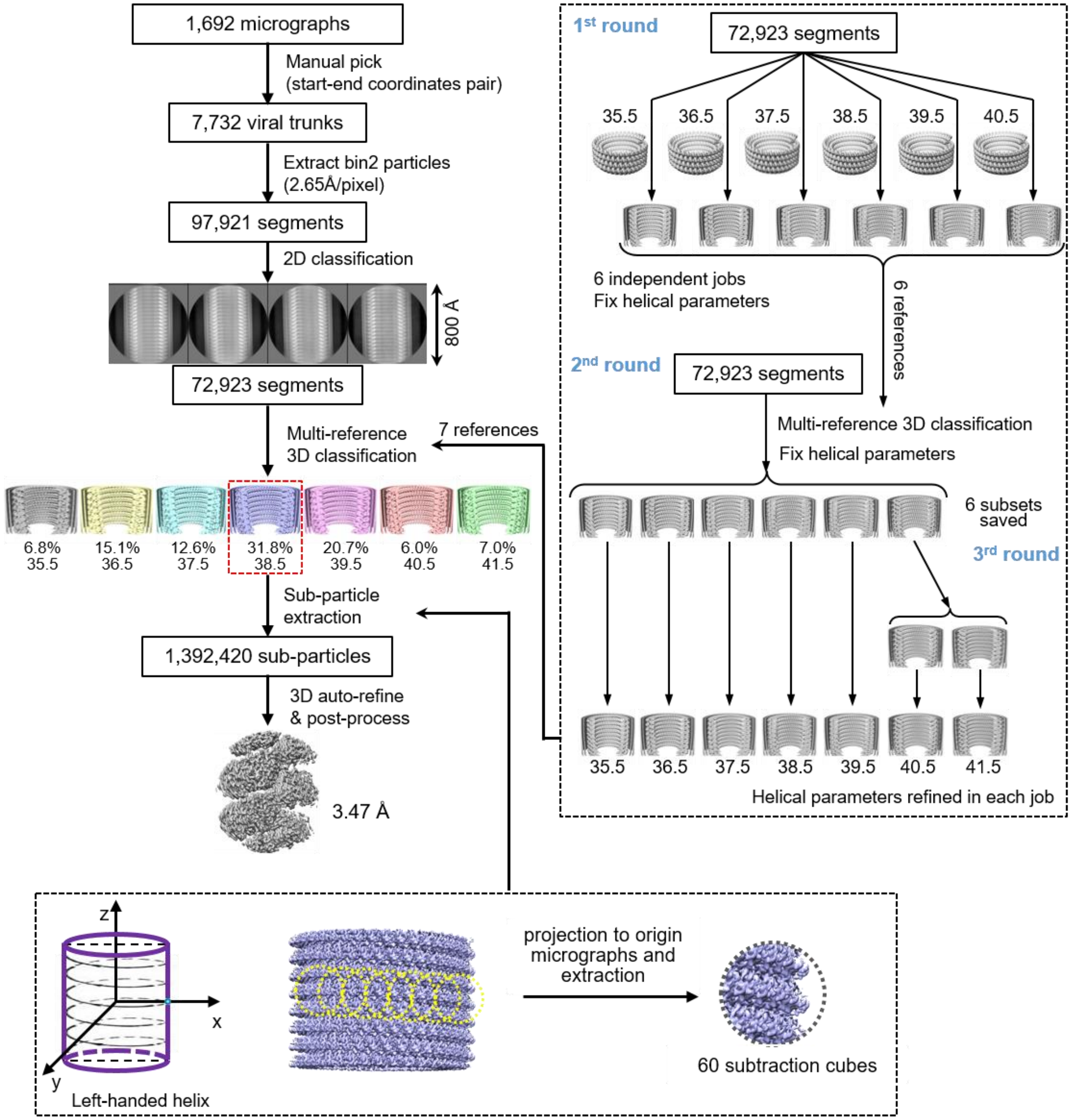


**Extended Data Fig. 7 | Data processing workflow for helical sub-particle reconstruction.**

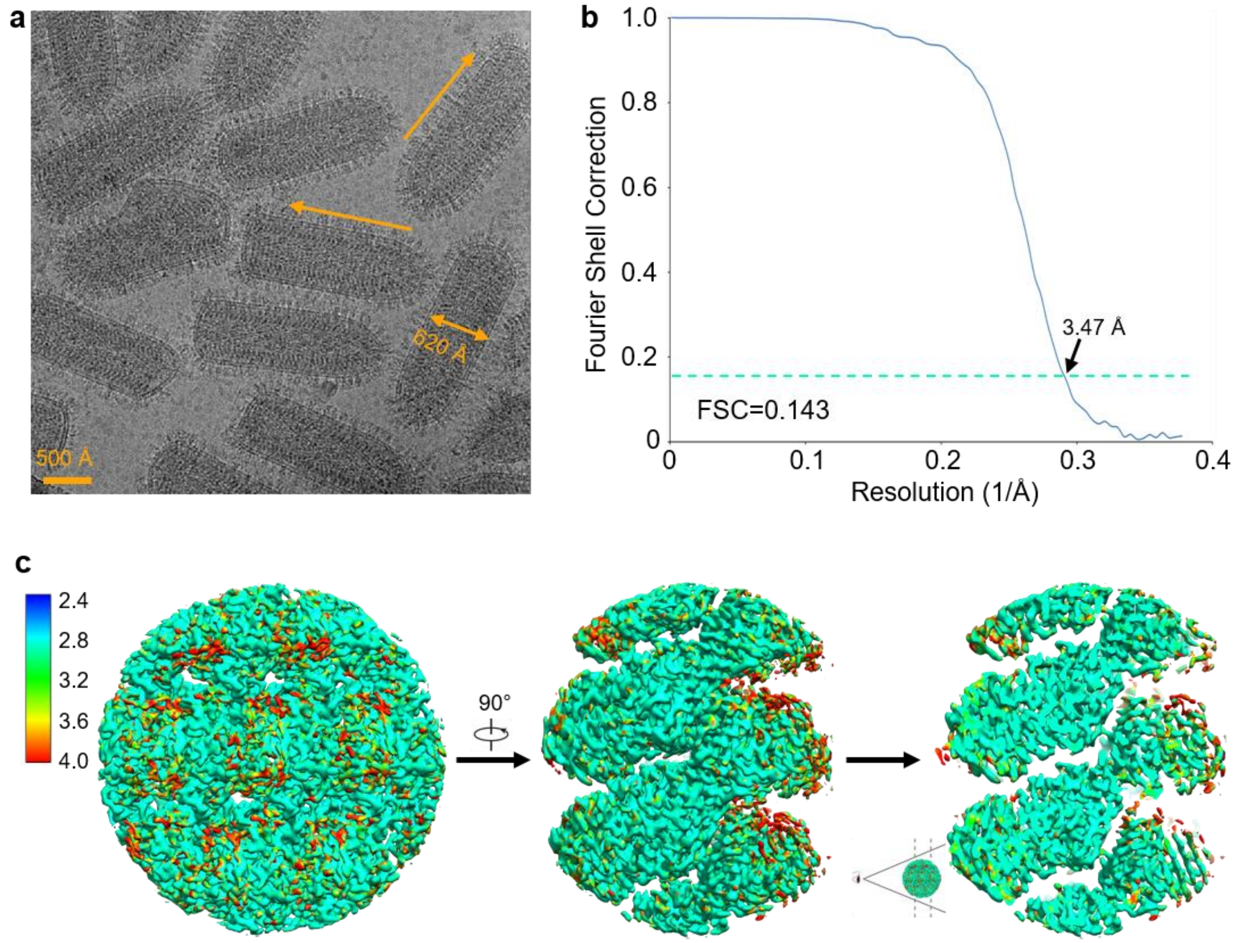


**Extended Data Fig. 8 | Near atomic resolution cryoEM analysis for the VSV trunk by helical sub-particle reconstruction. a:** A representative cryoEM micrograph of VSV virions. **b:** Global resolution evaluation based on "gold-standard" Fourier shell correction (FSC) coefficient as a function of spatial frequency generated by RELION, showing a resolution of 3.47 Å based on the 0.143 cut-off of FRC coefficient. **c:** Local resolution evaluation of the reconstructed sub-particle map.

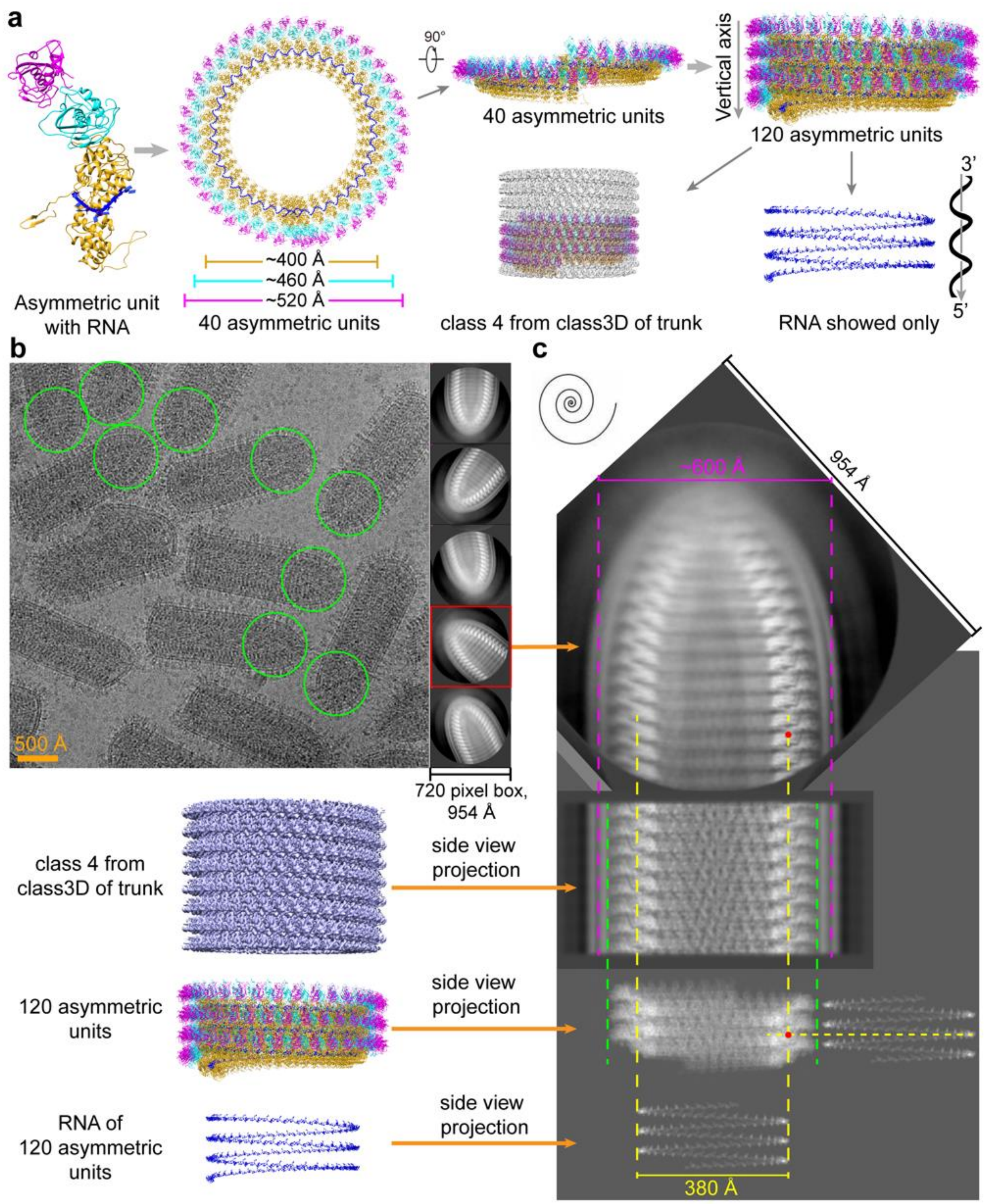


**Extended Data Fig. 9 | Preparation of the tip region for capsid reconstruction. a:** Helical turns obtained by applying helical symmetry (helical twist: -9.35°, helical rise: 1.3 Å) on the asymmetric units. The model of 120 asymmetric units fit perfectly in 3D class 4 from helical sub-particle reconstruction. **b:** A representative cryoEM micrograph of full VSV virions and 2D class averages of tips. The tips of virions are indicated by green circles. **c:** Alignment between the tip region and the trunk region. From top to bottom: 2D class of tips, projection of 3D class 4 from helical sub-particle reconstruction of trunk, projection of the 120 asymmetric unit model, and projection of RNA alone. The yellow dashed lines and red dots indicate the location of RNA inside 2D class of tips.

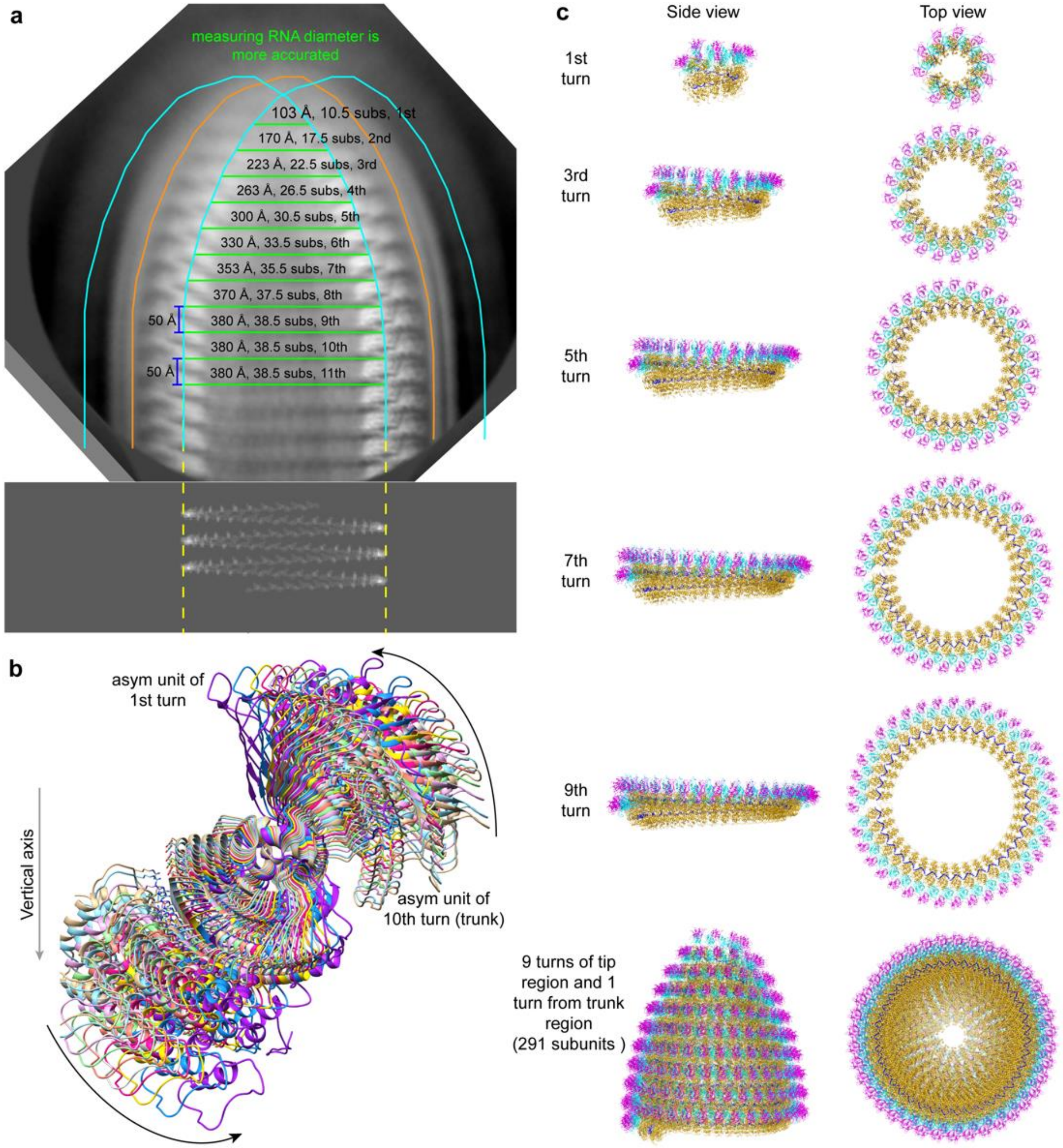


**Extended Data Fig. 10 | Tip region reconstruction. a:** Definition of the RNA strand edge in the tip region; the diameter of each RNA turn and their corresponding asymmetric unit (sub) numbers. **b:** The asymmetric unit model rotation of each turn in the tip region. **c:** Reconstructed model of turns and the composed model of the tip region.

#### Supplementary Table

| Subunit 1 | Subunit 2 | Interface area (Å²) | $\Delta^i G$ (kcal/mol) |
|---|---|---|---|
| $N_U$ | $N_{U+1}$ | 2551.6 | -20.8 |
| $N_{U-1}$ | $N_{U+1}$ | 321.4 | -5.4 |
| $N_U$ | $N_L$ | 29.5 | 0.1 |
| $N_U$ | $N_{L+1}$ | 180.8 | 0.5 |
| $IM_L$ | $N_U$ | 451.8 | 0 |
| **$IM_U$** | **$N_U$** | 398.9 | -4.4 |
| $IM_U$ | $N_{U+1}$ | 229.1 | -2.3 |
| $IM_U$ | $IM_L$ | 413.2 | -4.3 |
| **$IM_U$** | **$OM_U$** | 884.2 | -8.5 |
| $IM_U$ | $OM_{U-1}$ | 689.5 | -2.9 |

**Supplementary Table 1. Interface area and $\Delta^i G$ between subunits analyzed by PISA software.**

| Class number | Helical twist (°) | Helical rise (Å) | Subunits number per turn | Rise per turn (Å) |
|---|---|---|---|---|
| 1 | -10.14 | 1.42 | 35.50 | 50.41 |
| 2 | -9.86 | 1.38 | 36.51 | 50.38 |
| 3 | -9.60 | 1.34 | 37.50 | 50.25 |
| 4 | -9.35 | 1.30 | 38.50 | 50.05 |
| 5 | -9.12 | 1.27 | 39.47 | 50.13 |
| 6 | -8.89 | 1.24 | 40.49 | 50.21 |
| 7 | -8.68 | 1.21 | 41.47 | 50.18 |

**Supplementary Table 2. Helical parameters of 7 classes from 3D classification.**

| | VSV virion trunk |
|---|---|
| **Data collection** | |
| EM equipment | Titan Krios |
| Voltage (KV) | 300 |
| Detector | K2 summit |
| Nominal Magnification | ×105,000 |
| Pixel size (Å) | 1.325 |
| Total electron dose ($e^-/Å^2$) | 50 |
| Frame exposure time (s) | 0.2 |
| No. of movie frames | 45 |
| Defocus range (μm) | -1.5 ~ -3.0 |
| **Reconstruction** | |
| No. of micrographs collected | 2,008 |
| No. of micrographs used | 1,692 |
| Number of particles | 1,392,420 |
| Map sharpening B-factors ($Å^2$) | -170 |
| Resolution FSC 0.143 (Å) | 3.47 |
| Symmetry for final map | C1 |
| **Atomic model** | |
| No. of atoms | 41,417 |
| No. of protein residues | 5,022 |
| No. of nucleotide | 68 |
| Map CC | 0.78 |
| R.m.s deviations | |
| Bonds length (Å) | 0.003 |
| Bonds angle (°) | 0.633 |
| Ramachandran plot statistics (%) | |
| Preferred | 93.48 |
| Allowed | 6.52 |
| Outlier | 0 |
| Rotamers outliers (%) | 0.09 |
| C-beta deviations | 0 |
| MolProbity score | 2.01 |
| Clash score | 11.74 |
| PDB code | |
| EMDB code | |

**Supplementary Table 3. CryoEM data collection, refinement and validation statistics.**

| Turn | RNA cycle diameter (Å) | Subunit/turn | Helical twist (°) | Helical rise (Å) | Subunit rotation angle |
|---|---|---|---|---|---|
| 1st | 103 | 10.5 | -34.29 | 4.76 | --- (40 given) |
| 2nd | 170 | 17.5 | -20.57 | 2.86 | 31 |
| 3rd | 223 | 22.5 | -16 | 2.22 | 25 |
| 4th | 263 | 26.5 | -13.58 | 1.89 | 21.25 |
| 5th | 300 | 30.5 | -11.8 | 1.64 | 18.75 |
| 6th | 330 | 33.5 | -10.75 | 1.49 | 15 |
| 7th | 353 | 35.5 | -10.14 | 1.41 | 11.25 |
| 8th | 370 | 37.5 | -9.6 | 1.33 | 7.5 |
| 9th | 380 | 38.5 | -9.35 | 1.30 | 2.75 |
| 10th (first turn of trunk) | 380 | 38.5 | -9.35 | 1.30 | 0 |

**Supplementary Table 4. Calculated parameters of each turn in tip region.**

#### Supplementary Video Legends

**Supplementary Video 1 | Surface view of the sub-particle reconstruction of the VSV trunk.** The full density map is displayed as gray, followed by colored segmentations for intact OM (magenta), IM (cyan) and N (orange) with associated RNA (blue). Related to Figure 1.

**Supplementary Video 2 | Ribbon diagram of the atomic model built into the sub-particle reconstruction of the VSV trunk.** The model contains 5 OM (magenta) subunits, 7 IM (cyan) subunits and 7 N (orange) subunits and two RNA (blue) fragments. Related to Figure 2.

**Supplementary Video 3 | Slicing through a representative tomogram.**

**Supplementary Video 4 | The reconstructed entire VSV virion.** 360° panorama of entire VSV virion. N, IM, OM, RNA, L, P and G are colored in goldenrod, cyan, magenta, blue, red, black and green, respectively. Membrane is colored in gray.

**Supplementary Video 5 | Ribbon diagram of the VSV capsid.** Full capsid, partial layers hidden capsid and RNA alone were displayed.
